# Downy mildew effector HaRxL106 interacts with the transcription factor BIM1 altering plant growth, BR signaling and susceptibility to pathogens

**DOI:** 10.1101/2024.09.09.612066

**Authors:** María Florencia Bogino, Juan Marcos Lapegna Senz, Lucille Tihomirova Kourdova, Nicolás Tamagnone, Andres Romanowski, Lennart Wirthmueller, Georgina Fabro

## Abstract

*Hyaloperonospora arabidopsidis* (*Hpa*) is an oomycete pathogen that causes downy mildew disease on Arabidopsis. This obligate biotroph manipulates the homeostasis of its host plant by secreting numerous effector proteins, among which are the RxLR-effectors. Identifying the host targets of effectors and understanding how their manipulation facilitates colonization of plants is key to improve plant resistance to pathogens. Here we characterize the interaction between the RxLR effector HaRxL106 and BIM1, an Arabidopsis transcription factor (TF) involved in Brassinosteroid (BR) signaling. We report that HaRxL106 interacts with BIM1 *in vitro* and *in planta*. BIM1 is required by the effector to increase the host plant susceptibility to (hemi)biotrophic pathogens, and thus can be regarded as a susceptibility factor. Mechanistically, HaRxL106 requires BIM1 to induce the transcriptional activation of BR-responsive genes and cause alterations in plant growth patterns that phenocopy the shade avoidance syndrome. Our results support previous observations of antagonistic interactions between activation of BR signaling and suppression of plant immune responses and reveal that BIM1, a new player in this crosstalk, is manipulated by the pathogenic effector HaRxL106.

**SIGNIFICANCE STATEMENT:** Our study reveals that the Arabidopsis transcription factor BIM1 plays a key role as a susceptibility factor that is manipulated by the oomycete effector HaRxL106 to induce the Brassinosteroid (BR) signaling pathway and suppress plant immunity. HaRxL106 induces BIM1-dependent expression of BR-responsive genes which affects normal plant growth generating a shade avoidance-like syndrome that impacts negatively on plant defenses against bacterial and oomycete pathogens.

## INTRODUCTION

Plants have evolved complex mechanisms to resist the attack of multiple types of pathogens, but the energetic cost of inducing defenses in a resource-limiting environment can affect their growth and productivity (Belkhadir *et al.*, 2012; Huot *et al.*, 2014; Han and Kahmann, 2019). Conversely, certain plant organs or structures, either young, actively growing or nutritionally rich are considered more susceptible to colonization by pathogens (Wang and Wang, 2014, Lacaze and Joly, 2020). Plant defense mechanisms rely on cell surface and intracellular systems that detect specific pathogen-derived molecules (Dodds *et al.*, 2023). Cell-surface receptors can recognize compounds of microbial origin, called P/MAMPs (pathogen/microbe associated molecular patterns), like fungal chitin, bacterial flagellin or oomycete cell wall constituents. These receptors can also sense endogenous molecules derived from plant tissue damage. Recognized molecules are collectively named elicitors (Dodds *et al.*, 2023). The activation of cell surface receptors initiates a complex signaling cascade that includes Ca^2+^ increases in the cytosol, bursts of reactive oxygen species (ROS) at the plasma membrane and organelles, the activation of several mitogen-activated protein kinases and a wide transcriptional reprogramming leading to production of the defense hormones jasmonic acid (JA), ethylene and salicylic acid (SA). This response, called PTI (for Pattern-Triggered Immunity) usually provides a broad-spectrum resistance to a wide range of pathogens (DeFalco and Zipfel, 2021; Ngou *et al.*, 2022).

As a countermeasure, adapted pathogens, including oomycetes, secrete molecules called ‘effectors’ that interfere and rewire numerous plant metabolic pathways (Win *et al.*, 2012; Boevink *et al.*, 2020; He *et al.*, 2020). Oomycete pathogens produce multiple types of effectors, among them, the RxLR (Arg-x-Leu-Arg) family (Rehmany *et al.*, 2005; Win *et al.*, 2012). These effectors are secreted and translocated, probably via clathrin-mediated endocytosis (Wang *et al.*, 2023), into the cytoplasm of host cells. RxLR effectors redundantly manipulate multiple components of plant host cells, often referred to as ‘targets’ (He *et al.*, 2020; Fabro, 2022). Many host targets are directly involved in defense signaling while others are collaterally required for nutrient acquisition, delivery of pathogenicity factors or generation of adequate reproductive niches and are essential to maintain compatibility during invasion (Wang *et al.* 2019; He *et al.*, 2020; Judelson and Ah-Fong, 2019; Boevink *et al.* 2020).

Effectors can interfere with plant metabolic pathways that regulate plant growth and development. Both processes depend on photosynthesis-derived energetic resources which are also used to sustain plant defenses. The compromise of resource investment in one or the other process is known as the growth-defense tradeoff (Huot *et al.*, 2014; Züst and Agrawal, 2017; Han and Kahman, 2019). Although healthy plants, without environmental limitations, manage to balance active growth with an effective level of immunity, under sustained pathogen attack, plants destinate more energy and nutrients towards defense, leading to a reduced growth and alterations in development and reproduction (Paik *et al.*, 2017). For example, it has been shown that the activation of PTI and SA-signaling leads to growth inhibition (Navarro *et al.* 2006, Van Butselaar and van den Ackerveken, 2020). Conversely, numerous hormones involved in growth and development, such as auxins, gibberellins, cytokinins and brassinosteroids (BR), are able to modulate plant defenses (Robert-Seilaniantz *et al.* 2011; Igarashi *et al.*, 2012; Albretch *et al.*, 2012, Belkhadir *et al.*, 2012; Mosher and Kemmerling, 2013). Generally, rewiring of growth hormone signaling pathways by effectors results in a reduction in the level of plant defenses (Hahn and Kahmann, 2019; Dong and Ma, 2021; Figueroa *et al.*, 2021; Sperschneider and Dodds, 2022). BRs in particular, which are involved in cell elongation, vascular system differentiation, flowering time, photomorphogenesis, and other processes (Belkhadir and Jallais., 2015), play a central role in defenses against oomycete pathogens (Turnbull *et al.*, 2017, 2019).

The BR signaling pathway is activated upon perception of different BR hormones by the receptor BRASSINOSTEROID INSENSITIVE 1 (BRI1). This initiates a phosphorylation cascade involving the GSK3-like kinase BRASSINOSTEROID INSENSITIVE 2 (BIN2) that controls signal transduction via two major transcription factors: BRASSINAZOLE-RESISTANT 1 (BRZ1) and BR-INSENSITIVE-EMS-SUPPRESSOR 1 (BES1)/BZR2 (Yin, *et al.*, 2002; Belkhadir *et al.*, 2006; Wang *et al.*, 2014). BZR1 binds to BRRE motifs in the promoters of several BR-responsive genes, while BES1 forms heterodimers with another bHLH transcription factor (TF) named BIM1 (BES1-interacting MYC2-Like1, At5g08130) and together bind to E-boxes (CACGTG) in the promoters of numerous genes (Yin *et al.*, 2005; Zhu *et al.* 2013). BIM1 positively modulates hypocotyl elongation and the shade avoidance syndrome in Arabidopsis seedlings via its interaction with the transcription factor PHYTOCHROME RAPIDLY REGULATED1 (PAR1) (Yin *et al.*, 2005; Cifuentes-Esquivel *et al.* 2013).

Activation of BR signaling can either increase or antagonize defense responses against pathogens, depending on the pathosystem considered, and if the activation is either endogenous or via an exogenous application (Albretch *et al.*, 2012; Belkhadir *et al.*, 2012; Lozano-Durán *et al.*, 2013; Malinovsky *et al.*, 2014). In Arabidopsis, the activation of BR signaling suppresses PTI-induced transcriptional responses mainly through the activity of BRZ1, which regulates the activation of WRKY transcription factors that negatively impact on PTI-related gene expression (Lozano-Durán *et al.*, 2013). Similarly, BRZ1 also activates the bHLH transcription factor HBI1, that negatively regulates the expression of PTI-marker genes (Fan *et al.*, 2014; Malinovsky *et al.*, 2014). On the contrary, treatment with brassinazole that inhibits BR biosynthesis, increases the ROS burst in response to PAMPs (Lozano-Durán *et al.*, 2013). In fact, there are numerous components shared between the BR and PTI signaling, that have been proposed to modulate the balance between growth and defenses (Chinchilla *et al.* 2009; Lin *et al.*, 2013; Fan *et al.*, 2014; Kang *et al.*, 2015; Bücherl *et al.*, 2017;Ortiz-Morea *et al.*, 2020). Interestingly, regarding plant-oomycete interactions, transgenic plants overexpressing BRI1 show increased susceptibility to the biotrophic oomycete *Hyaloperonospora arabidopsidis* (Hpa) (Belkhadir *et al.* 2012). In line with this, it has been reported that the overexpression of the oomycete RxLR effector AVR2 from *Phytophthora infestans* induces BR signaling in potato which leads to a suppression of PTI-dependent defenses (Turnbull *et al.*, 2017, 2019).

By using the model pathosystem Arabidopsis-Hpa we here report the role of a new target of the effector HaRxL106, the transcription factor BIM1, in the regulation of Arabidopsis growth-defense tradeoff. It was previously described that HaRxL106 is actively transported into the plant cell nucleus and inhibits PAMP triggered ROS burst as well as SA-dependent gene expression (Fabro *et al.*, 2011; Wirthmueller *et al.* 2015 and 2018) enhancing susceptibility to compatible Hpa isolate NoCo2 (Wirthmueller *et al.* 2018) as well as to the virulent bacteria *Pseudomonas syringae* (Pst) (Fabro *et al.*, 2011). Moreover, HaRxL106 stable expression in Arabidopsis (HaRxL106-OE) generates an altered growth phenotype, with elongated petioles, and narrow, curved down leaves that is reminiscent of plants displaying constitutive shade avoidance syndrome (SAS) (Fabro *et al.* 2011, Wirthmueller *et al.*, 2018). The C-terminal 58 amino acids of HaRxL106 are sufficient and required to suppress host defenses and induce the SAS-like phenotype when expressed as transgene (Wirthmueller *et al.* 2018). We here describe our recent findings indicating that the SAS-like developmental phenotypes may be explained by the interaction of HaRxL106 with BIM1. HaRxL106-BIM1 interaction is direct and occurs in plant nuclei. Our results indicate that HaRxL106 affects the activity of BIM1 as a regulator of BR-triggered gene expression, and that BIM1 is required by the effector to confer enhanced susceptibility and alter plant growth pattern.

## RESULTS

### HaRxL106 interacts with BIM1

It was previously demonstrated that the SAS-like phenotype of HaRxL106-OEs depends on the interaction of the effector with the transcriptional regulator RADICAL-INDUCED CELL DEATH1 (RCD1) but due to the multiple pleiotropic effects that the loss of RCD1 causes in Arabidopsis, the underlying mechanism remains elusive (Wirthmueller *et al.*, 2018). Looking for other HaRxL106 interactors whose function can be related to the above-mentioned phenotype we took advantage of a preliminary yeast-two-hybrid (Y2H) screen performed in Dr. Jonathan D.G. Jones laboratory (personal communication) which suggested that HaRxL106 interacted with BIM1. We also observed that the E-box motif (CACGTG) was enriched in the promoters of genes that showed upregulated expression in HaRxL016-OE Arabidopsis lines (Wirthmueller *et al.* 2018, and Table S6 therein), either in non-treated or mock treated plants as well as in those infected with Pst. Therefore, we set up a directed Y2H experiment to verify the HaRxL106-BIM1 interaction and determine which parts of HaRxL106 are required for it. We thus evaluated BIM1 interaction with the full length (FL) version of the effector, with the last 58 amino acids of its carboxy-terminal domain (c58) and with the amino terminal part of HaRxL106 (ΔC), lacking the last 58 amino acids. As it can be observed in **Figure 1A**, the full length (HaRxL106-FL) and the C-terminal domain of HaRxL106 (HaRxL106-c58) interact with BIM1 while the N-terminal domain (HaRxL106-ΔC) does not interact. The negative controls did not show autoactivation of neither BIM1 nor HaRXL106. To investigate if the interaction between HaRxL106 and BIM1 was direct, as the Y2H result suggested, we performed a pull-down experiment with recombinant full-length versions of these proteins expressed in *E. coli*. We affinity purified BIM1 tagged with MBP (MBP-BIM1, Liang *et al.*, 2018) and HIS-HaRxL106 (Wirthmueller *et al.*, 2015). As it can be observed in **Supplementary Figure S1**, MBP-BIM1 bound to an amylose resin pulls down HIS-HaRxL106. This interaction does not occur with MBP alone.

**Figure 1:**
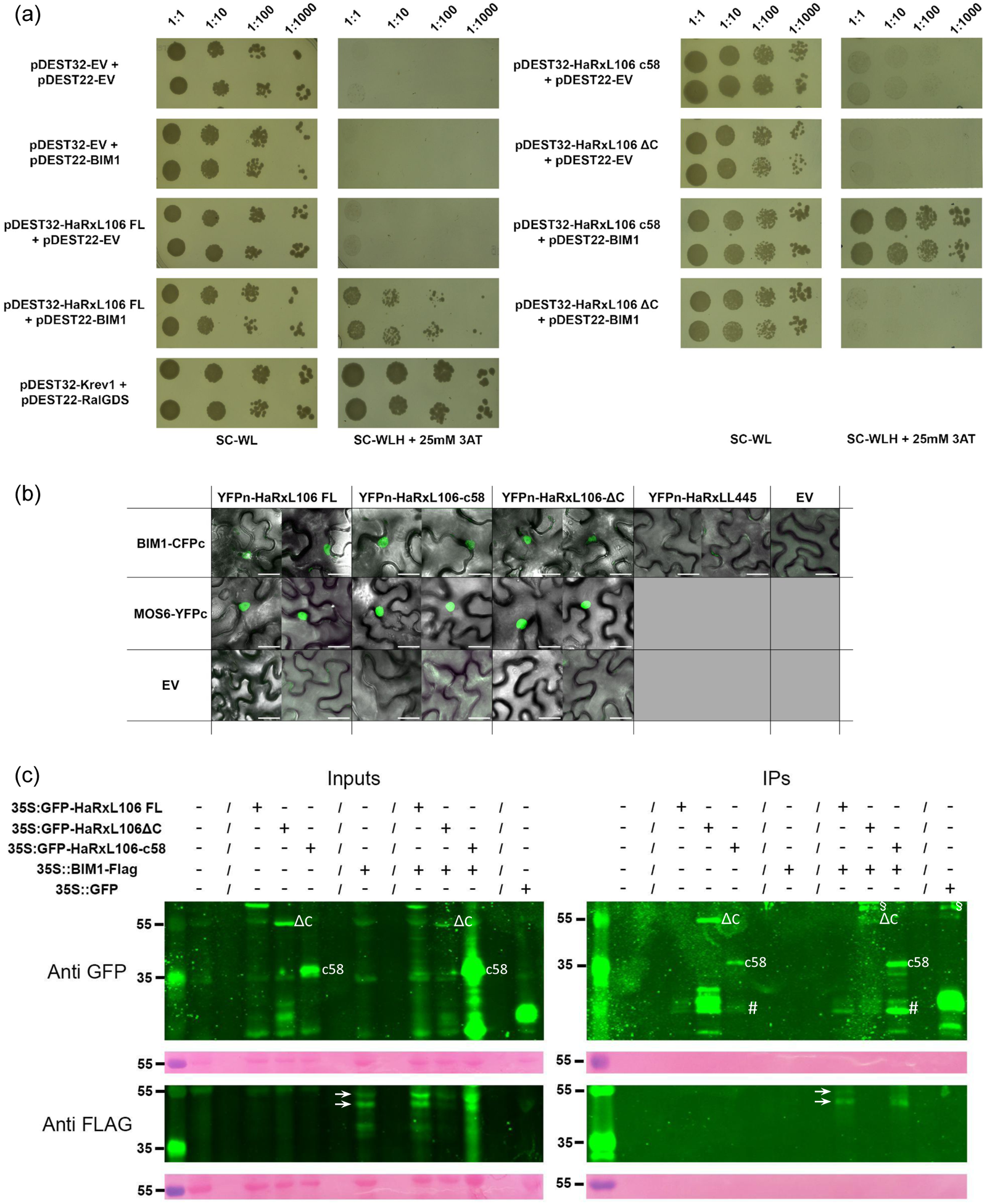
HaRxL106 directly interacts with BIM1. **a.** Y2H assay using HaRxL106 effector full length (FL) sequence, and its carboxyl (HaRXL106-c58) and amino-terminal (HaRxL106-ΔC) domains. pDEST32: bait vector. pDEST22: prey vector. SC-WL: medium Synthetic Complete for yeasts without Tryptophan and Leucine. SC-WLH + 25 mM 3AT: SC-WL and without Histidine, supplemented with 25mM 3-Aminotriazole. pDEST32-Krev1+pDEST22-RalGDS: positive interaction control. **b**. Bi-molecular fluorescence complementation assay. Confocal microscopy images representative of the abaxial epidermis of *N. benthamiana* leaves co-expressing the indicated constructs. Scale bars represent 20 µm. **c**. Co-Immunoprecipitation of GFP-HaRxL106 FL or GFP-HaRxL106-c58 with BIM1-FLAG in Arabidopsis. Left panels: SDS-PAGE of total protein extracts (inputs), revealed with anti-GFP (upper panel) or anti-FLAG antibodies (lower panel). Right panels: proteins immunoprecipitated with Magne-Halo-anti-GFP, revealed with anti-GFP (upper panel) or anti-FLAG antibodies (lower panel). Ponceau S staining shows RUBisCo loading. (/) indicates an empty well, white arrows point to BIM1-FLAG degradation bands. (§) nonspecific anti-GFP antibody binding, (#) GFP-HaRxL106 degradation bands. Line 2 on the left of all blots, next to the MW weight marker is Col-0 wild type protein extract.

It was previously reported that HaRxL106 is transported into the nucleus by several IMPORTIN-α isoforms that bind to its C-terminal nuclear localization sequence (NLS) (Wirthmueller *et al.* 2015), and also that BIM1 has a predicted NLS (Hooper *et al.*, 2017). Given that the role of BIM1 as a transcription factor depends on its nuclear localization, we performed Bi-molecular fluorescence complementation (BiFC) assays to verify if the interaction with HaRxL106 takes place in the plant nucleus. For this, we used the three versions of the effector (HaRxL106 FL, HaRxL106 ΔC and HaRxL106 c58) to express them transiently in leaves of *Nicotiana benthamiana* (**Figure 1B**). The variants of the effector were tagged at their N-terminus with the YFPn protein fragment to avoid interference with the carboxy-terminal effector domain. BIM1 was tagged at its C-terminus with the CFPc fragment as it was previously reported that this tag orientation did not interfere with its activity (Yin *et al.* 2005). As positive control we used importin-a3/MODIFIER OF SNC1 6 (MOS6), a known interactor of HaRxL106, tagged at its C-terminus with YFPc (Wirthmueller *et al.*, 2015). As a negative control we used another nuclear Hpa effector (HaRxLL445) (Caillaud *et al.*, 2012). As can be observed in **Figure 1B**, HaRxL106-FL and HaRxL106-c58 interacted with BIM1 in the nucleus of *N. benthamiana* epidermal cells. In contrast to the Y2H results, BiFC also showed interaction between HaRxL106-ΔC and BIM1 (**Figure 1B**). This might indicate a lower level of stringency in BiFC compared to theY2H assay. On the other hand, BIM1 did not show fluorescence complementation with the negative control HaRxLL445.

BIM1 is part of a protein family that has two other members, BIM2 (At1g69010) and BIM3 (At5g38860). The major difference between them is that BIM1 has an amino-terminal domain (N-term) that is absent in BIM2 and BIM3 (Yin *et al.*, 2005). To assess if HaRxL106 also interacts with these nuclear proteins, we performed a BiFC assay with BIM2 and BIM3 (**Supplementary Figure S2**). We observed that both interacted with HaRxL106-FL, suggesting that the N-term of BIM1 is not the sole domain providing the interaction interface for HaRxL106. This result is further substantiated by a Y2H assay between the N-term of BIM1 (amino acids 1 to 278) and the C-terminus of HaRxL106 (HaRxL106-c58) that, if at all, showed a very weak interaction (**Supplementary Figure S3**). In contrast, the interaction between the N-term of BIM1 and the FL version of the effector could be detected in this assay. Although the combined *in planta* and Y2H results do not define an unambiguous mode of interaction, our data are consistent with a model in which the N-terminal part of HaRxL106 generally binds to the part that is shared by BIM1 and BIM2. Based on the BiFC results (**Figure 1B, Supplementary Figure S2**), the C-terminal 58 amino acids of the effector provide additional specificity for the association with BIM1 although the unique N-terminal part of BIM1 on its own is not sufficient for binding HaRxL106 in Y2H (**Supplementary Figure S3**).

Given the caveats that BiFC techniques have (Ohad *et al.* 2007), to further verify the interaction between HaRxL106 and BIM1 and clearly determine the parts of the effector that were necessary and sufficient for the interaction *in planta*, we performed co-immunoprecipitation assays (CoIP) by transiently expressing GFP-HaRxL106-FL and GFP-HaRxL106-c58 together with BIM1-FLAG (Yin *et al.*, 2005) in leaves of *N. benthamiana* (**Supplementary Figure S4 A, B**). Although BIM1 undergoes partial degradation during purification from *N. benthamiana,* a portion of the protein can be co-immunoprecipitated by GFP-HaRxL106-FL (**Supplementary Figure S4A**) and also by the C-terminal domain of the effector (**Supplementary Figure S4B**). Another GFP-tagged nuclear Hpa effector (HaRxLL445) did not immunoprecipitate BIM1-FLAG (**Supplementary Figure S4A**).

Finally, to verify if the effector indeed interacted with BIM1 in Arabidopsis, we used transgenic plants generated by other authors (Liang *et al.*, 2018) that, as we verified, constitutively expressed BIM1-FLAG (**Supplementary Figure S5**). We stably transformed these transgenics to introduce the different domains of the effector, generating double transgenics that co-expressed BIM1-FLAG and GFP-HaRxL106-FL or ΔC or c58 to perform CoIP assays in Arabidopsis seedlings. We observed that BIM1-FLAG could be immunoprecipitated by GFP-HaRxL106-c58 as well as by HaRxL106-FL, even when the full-length version of the effector became rapidly degraded during the IP process (**Figure 1C**). Conversely, HaRxL106-ΔC did not interact with BIM1-FLAG (**Figure 1 C**). Thus, by using different techniques, our results strongly suggest that HaRxL106 physically interacts with the transcription factor BIM1. Consistent with the interaction model presented above, the 58 C-terminal amino acids of HaRxL106, that mediate suppression of plant defense and confer the SAS-like phenotype (Wirthmueller *et al.*, 2018), is also necessary and sufficient for the interaction with full-length BIM1 but not with its N-terminal domain alone or with the other Arabidopsis members of this transcription factor family, BIM2 and BIM3.

### HaRxL106 requires BIM1 to confer hyper-susceptibility to *Hpa* and *Pst*

To investigate if BIM1 influences Arabidopsis immunity towards Hpa, we first determined sporulation levels of Hpa race NoCo2 on *bim1* mutants (**Figure 2A**). For this we used a line with a T-DNA insertion in the 9^th^ exon of BIM1 (SALK_132178) (**Supplementary Figure S6 A, C, D**) and BIM1-FLAG overexpressing plants (**Figure 2A**). An independent, previously published *bim1* T-DNA line (SALK_085924, insertion in 1^st^ exon), did not harbor the reported T-DNA insertion and was therefore excluded from the analysis (see Materials and Methods section and **Supplementary Figure S6 B**). However, we were able to characterize the SALK_085924 *bim1* mutation in context of the *bim123* triple mutant including genetic complementation, see below. We also assessed the susceptibility of *bim2* and *bim3* mutants to Hpa (**Supplementary Figure S7A**). All three single mutants *bim1*, *bim2* and *bim3* showed decreased conidiospore production when compared to Col-0 wild type plants (**Figure 2A**, **Supplementary Figure S7A**). Hpa conidiation was also reduced in the triple mutant *bim123* (**Figure 2A, Supplementary Figure S7B**) indicating that loss of single or multiple BIM genes could either activate immune signaling or BIM proteins might act as susceptibility factors for Hpa. We observed a trend towards increased Hpa conidiation on BIM1-FLAG overexpressing plants, similar to the one observed in plants that overexpress the effector HaRxL106 although this was not always statistically significant for BIM1-FLAG over-expressors (**Figure 2A, Supplementary Figure S7B**). Complementation of the triple mutant *bim123* with BIM1-FLAG restored the wild type levels of susceptibility to Hpa (**Supplementary Figure S7B**), indicating that BIM1 on its own is sufficient to act as a ‘susceptibility factor’ which could explain the increased resistance observed in *bim1* plants. We note that our phenotyping results do not exclude possible additional functions of BIM2 and BIM3 as susceptibility factors. However, based on the specific interaction between the defense-manipulating C-terminal part of HaRxL106 and BIM1, we focus here on BIM1 as a possible virulence target of the effector.

**Figure 2:**
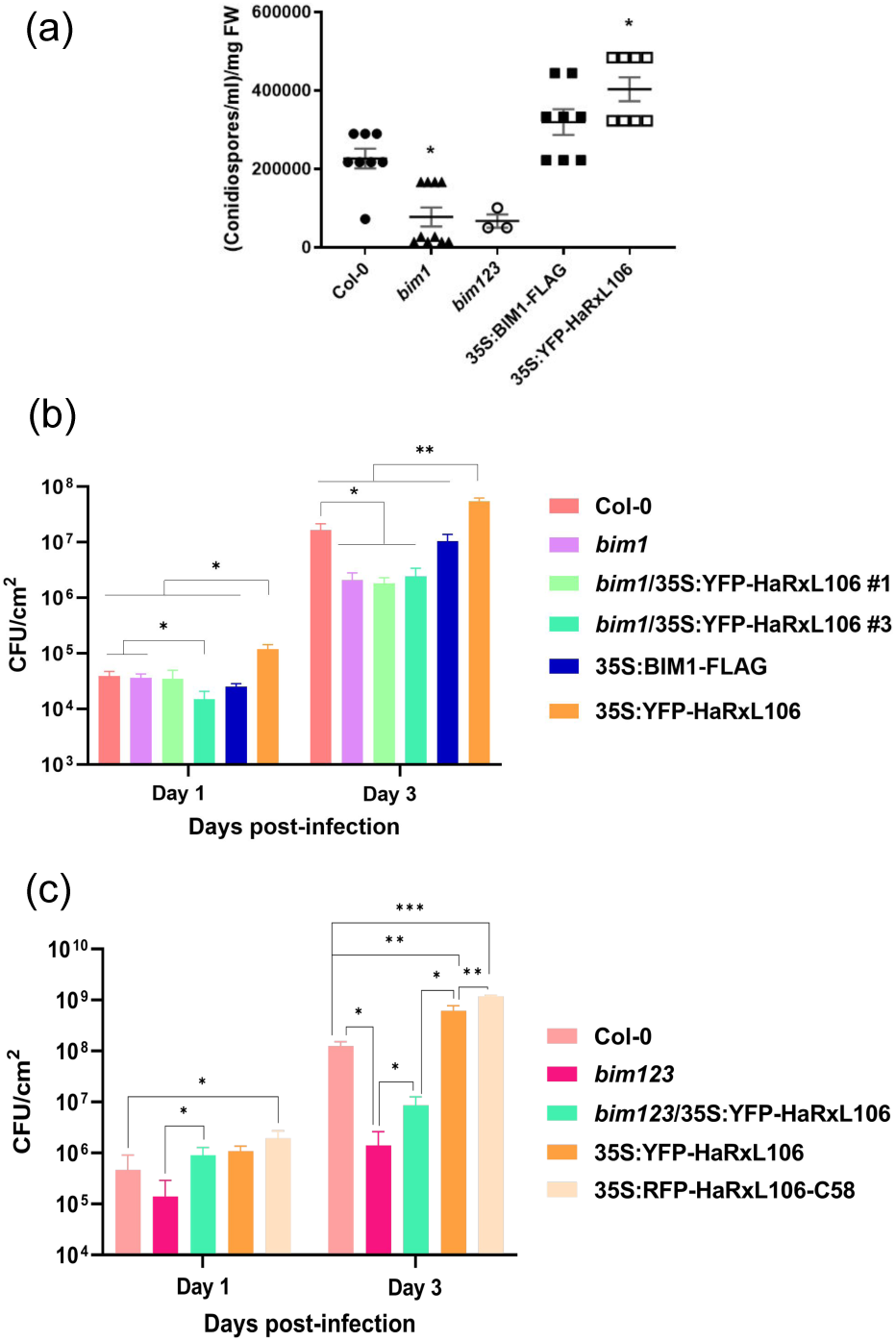
*bim1* and *bim123* mutants are less susceptible to *Hyaloperonospora arabidopsidis* (Hpa) and *Pseudomonas syringae* pv. tomato DC3000 (Pst) and BIM1 is necessary for the increased susceptibility to Pst of HaRxL106 overexpressing plants. **a.** Number of Hpa conidia produced by infected A. thaliana seedlings (n=50) of the indicated genotypes. The asterisks indicate significant differences compared to Col-0 according to a non-parametric ANOVA (Kruskal Wallis) with p<0.05. This experiment was repeated 6 times with similar results. **b-c.** Bacterial titers in colony-forming units per square centimeter of leaf (CFU/cm^2^) at 1 and 3 days after infection with Pst in the genotypes mentioned. Significant differences with Col-0 or between the genotypes indicated according to the t-test for averages differences with p<0.05 (*), p <0.01 (**) and p<0.001 (***) are indicated. n= 6 for B; n=3 for C. The experiments were repeated 3 times with similar results.

Given the fact that we had already observed that HaRxL106-OE lines, besides displaying hyper-susceptibility to Hpa, were also more susceptible to virulent strains of the bacterium Pst (Fabro *et al.* 2011), we decided to also assess the response of *bim1* mutant plants to Pst. This pathogen inoculation is easier to synchronize and its growth *in planta* is simpler to quantitate over time. We verified the previous observation and noted that plants expressing RFP-HaRxL106-C58 were the most susceptible (**Supplementary Figure S7C and Figure 2C**). We also observed that *bim1* plants supported lower bacterial growth by day 3 post inoculation (**Figure 2B**, **Supplementary Figure S7C**). Moreover, *bim123* triple mutants showed an even greater decrease in bacterial titers from day 1 that was maintained by day 3 post infection (**Figure 2C**, **Supplementary Figure S7C**). To investigate if BIM1 was also required for the enhanced bacterial susceptibility caused by the expression of HaRxL106 in wild type plants, we generated transgenics that expressed the effector in the *bim1* mutant background. We confirmed by qPCR and confocal microscopy that HaRxL106 was indeed expressed in the *bim1* background (**Supplementary Figure S8 A, B**). To assess the potential roles of BIM2 and BIM3 in HaRxL106-mediated bacterial susceptibility, we also generated HaRxL106 overexpression lines in the *bim23* double and *bim123* triple mutant backgrounds. As can be observed in **Figures 2B and 2C**, when the effector HaRxL106 is expressed in the absence of BIM1, it cannot confer enhanced susceptibility to Pst. Additionally, and in contrast to the phenotype observed for Hpa (**Supplementary Figure S7A**), BIM2 and BIM3 do not appear to be required for HaRxL106 to enhance Arabidopsis susceptibility to the bacteria (**Supplementary Figure S7D**). Thus, the enhanced susceptibility to both pathogens produced by the expression of HaRxL106 specifically requires BIM1, supporting the idea that this protein may act as a susceptibility factor that is required to establish a fully compatible interaction with Hpa and Pst.

### *bim1* mutants show altered PTI responses

The enhanced resistance of *bim1* to Hpa and Pst could either be due to some type of constitutive defenses activation (e.g. PTI), indicating that BIM1 plays a role as a negative regulator of plant defenses, or to the requirement of BIM1 as a susceptibility factor to establish a proper infection process (Fabro, 2022). To investigate this, we evaluated if *bim1* mutants showed signs of constitutive or enhanced PTI by monitoring callose deposition induced by the Pst *hrcC-* mutant strain (*Pseudomonas syringae* pv. tomato DC3000 *hrcC-*, Yuan and He, 1996), ROS burst induced by the bacterial PAMP flg22, and transcriptional activation of defense marker genes. As shown in **Figure 3A**, *bim1* does not show increased basal levels of callose deposition. Strikingly, while wild type plants responded with elevated callose deposition to the infection with Pst *hrcC-*, the *bim1* mutant was completely devoid of pathogen-induced callose deposits. Additionally, flg22-induced ROS levels in *bim1* mutants were 40% lower than those observed in wild type plants (**Figure 3B**). Regarding the expression levels of marker genes of the SA (*ICS1* and *PR1*) or JA (*PDF1.2*) defense pathways, their basal levels were either undetectable or highly variable between genotypes and experiments without a clear correlation with the presence/absence of BIM1 (**Supplementary Figure S9**). In line with this, the transcriptional expression of these marker genes upon Hpa infection showed no correlation with the susceptibility or resistance phenotypes we observed previously (**Supplementary Figure S9**, **Figure 2**). Thus, the enhanced resistance of *bim1* mutants to Hpa and Pst does not appear to be due to a direct effect of BIM1 as a negative regulator of plant defenses (PTI or SA/JA pathways). Conversely, the reduced susceptibility of *bim1* mutant plants might be an indirect consequence of impeded activation of the BR signaling pathway, which prevents alterations in plant growth and development that hijack resources that could be invested in plant defense, weakening plant cell walls, facilitating pathogen penetration and/or promoting metabolic changes that favor pathogen nutrition and reproduction. This could also explain why the reduced susceptibility is observed for two pathogens from different kingdoms and different lifestyles (Hpa, an oomycete obligate biotroph, Pst, a bacterial hemibiotroph).

**Figure 3:**
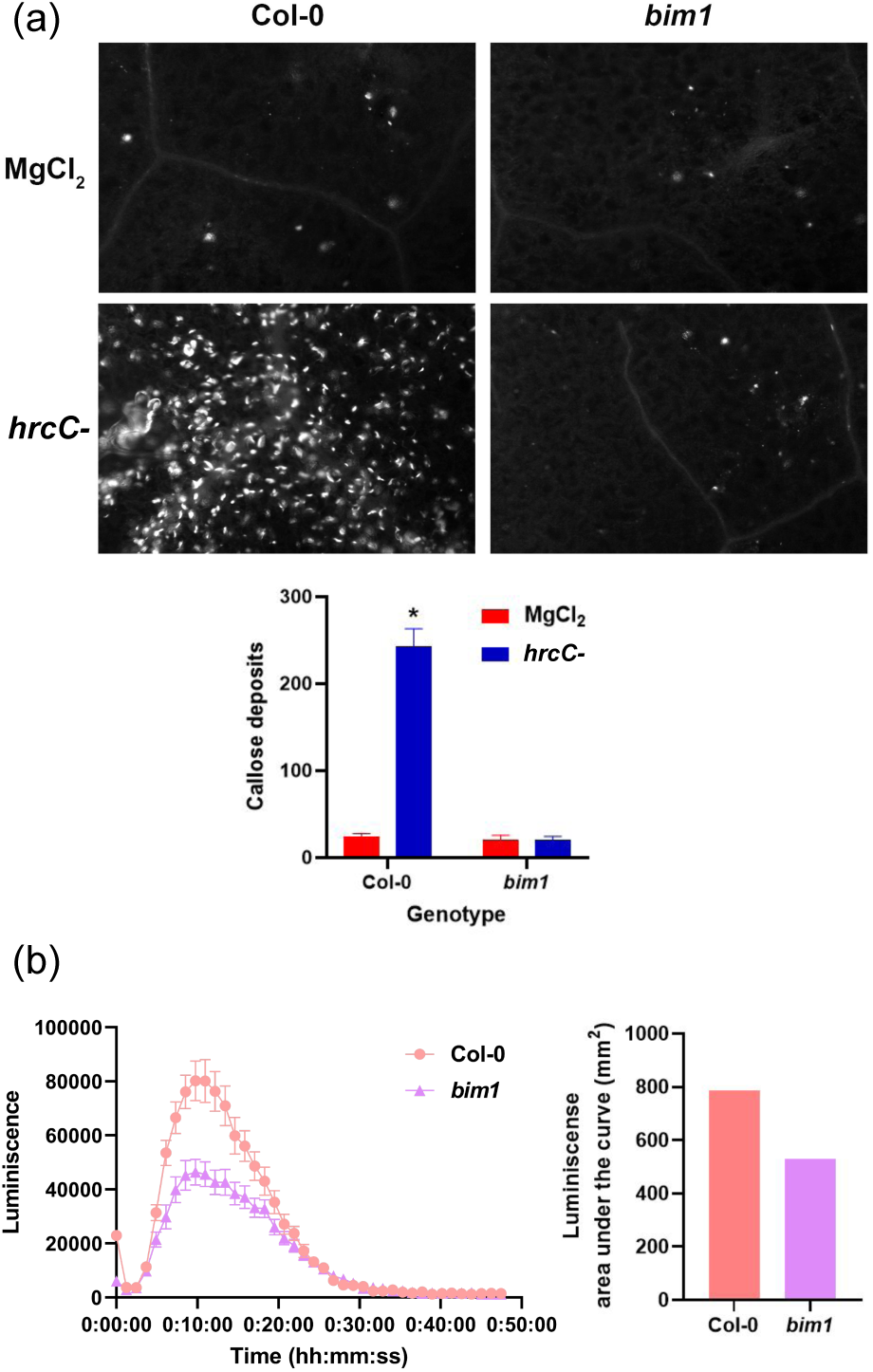
*bim1* mutants show altered PTI-responses. **a.** Callose deposits on the abaxial epidermis of leaves of the indicated genotypes treated with buffer (MgCl_2_ 10 mM) or infected with *Pseudomonas syringae* pv. tomato DC3000 (Pst) mutant hrcC-. The asterisk indicates significant differences with p<0.05 according to a non-parametric ANOVA (Kruskal Wallis), n=36. **b.** Kinetics of the oxidative burst (ROS) after treatment with flg22 of leaf discs (n=24) of the indicated genotypes. Measure of luminescence is expressed in Relative Luminescence Units (RLUs).

### BIM1 is necessary for HaRxL106 to cause SAS-like developmental phenotypes

As mentioned, expression of HaRxL106 as a transgene in the Col-0 background induces a constitutive shade avoidance syndrome (SAS)-like phenotype (Fabro *et al.* 2011, Wirthmueller *et al.*, 2018). It was previously demonstrated that this phenotype depends on the interaction of HaRxL106 with the transcriptional co-regulator RADICAL-INDUCED CELL DEATH1 (RCD1) but due to the multiple pleiotropic effects that the loss of RCD1 causes in Arabidopsis, the mechanism by which HaRxL106 triggers a SAS-like developmental phenotype remains elusive (Wirthmueller *et al.* 2018).

Thus, to determine whether BIM1 was required by HaRxL106 for the development of the SAS-like phenotype, we evaluated developmental parameters on *bim1* and *bim123* plants expressing the full length YFP-tagged version of HaRxL106 (35S:YFP-HaRxL106). Only one of the two independent lines, *bim1*/35S:YFP-HaRxL106 (line #3), had a slightly greater hypocotyl length than Col-0 and *bim1* seedlings (**Figure 4A**). However, as adult plants, none of the lines recapitulated the phenotype observed in 35S:YFP-HaRxL106 over-expressors in wild type Col-0 background (**Figure 4B**).

**Figure 4:**
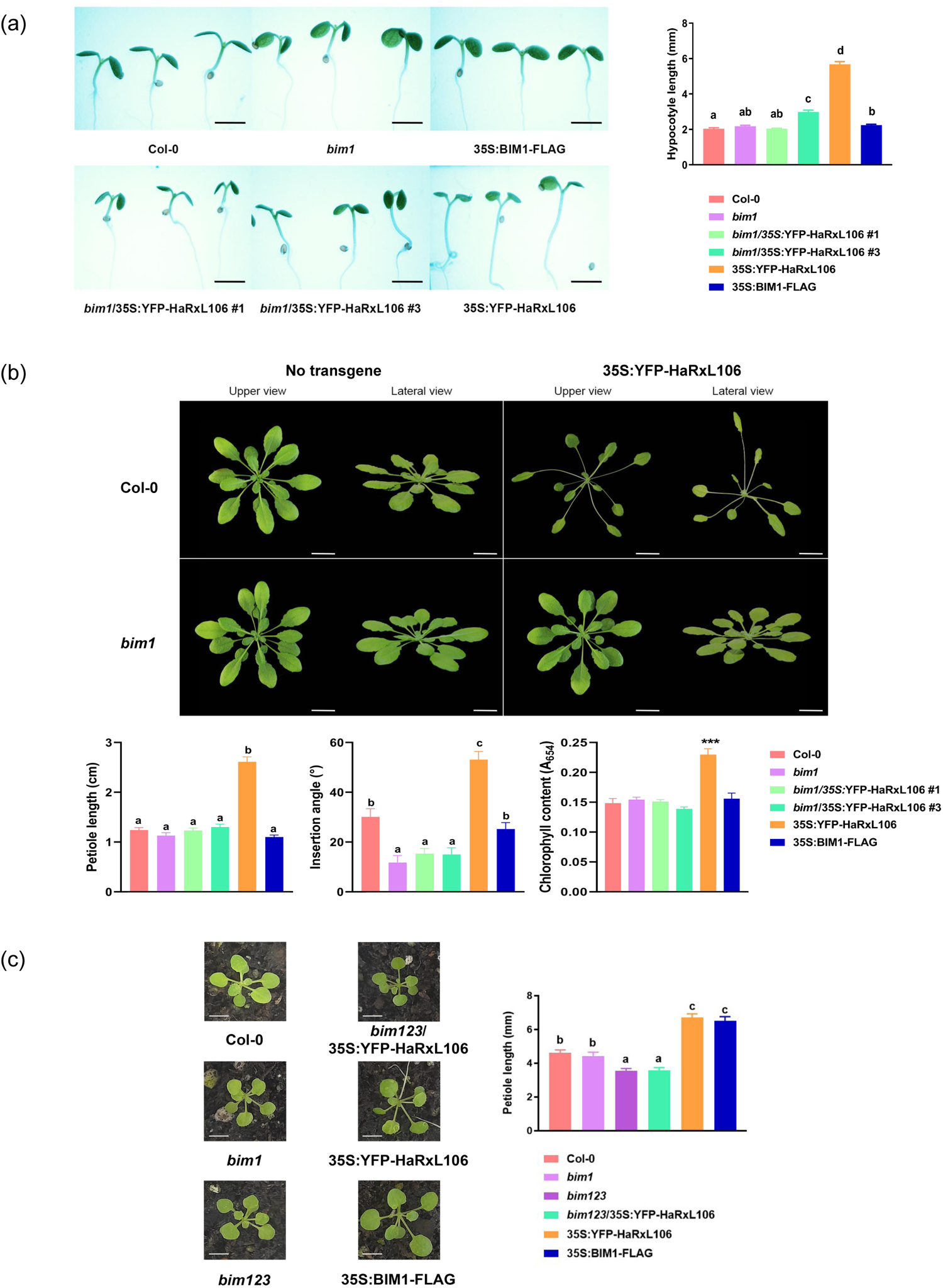
HaRxL106 expression in planta alters plant development in a BIM1-dependent manner. **a.** Representative images of 10 day-old seedlings of the indicated genotypes and quantification of their hypocotyl length. Different letters indicate significant differences with p<0.05 according to non-parametric ANOVA (Kruskal Wallis) n=26. Scale bars represent 2 mm. **b.** Representative images of two month-old adult plants overexpressing HaRxL106 in wild type (Col-0) or *bim1* mutant background, (Col-0/YFP-HaRxL106 #12 and *bim1*/YFP-HaRxL106 #3). Scale bars represent 2 cm. Quantification of petiole length, petiole insertion angle and chlorophyll content of adult plants of the genotypes (colour coded as in A) is shown. Different letters indicate significant differences according to non-parametric ANOVA (Kruskal Wallis, p<0.001); and asterisks according to a parametric ANOVA with Tukey post hoc test (p<0.001). c. Representative images of 3 week-old seedlings and petiole length quantification. Different letters indicate significant differences with p<0.05 according to non-parametric ANOVA (Kruskal Wallis). Scale bars represent 5 mm.

Col-0 constitutively expressing HaRxL106-FL (35S:YFP-HaRxL106) also shows leaves with elongated petioles, which are inserted into the rosette at larger angles (leaf hyponasty). Thus, we also evaluated the petiole length and leaf insertion angle in young (3-week-old) or adult (2-months-old) *bim1* mutants comparing them with plants expressing HaRxL106 (**Figure 4B** bar graphs and **Figure 4C**). *bim1* mutants showed no significant differences in petiole length regarding wild type plants either at 2 months (**Figure 4B**) or 3 weeks (**Figure 4C**) of age. We could observe that 2-month-old *bim1* rosettes were flatten with a smaller insertion angle than wild type plants. The plants that expressed the oomycete effector also showed a higher chlorophyll content per square centimeter of leaf lamina (**Figure 4 B,** far right graph). On the contrary, *bim1* mutant plants were similar to Col-0 plants in their chlorophyll content. Young BIM1-FLAG plants (3-weeks-old) showed elongated petioles, similar to plants expressing HaRxL106 (**Figure 4 C**) but this phenotype disappeared over time. Two-month old BIM1-FLAG plants were identical to Col-0 (**Figure 4B**, petiole length graph and **Supplementary Figure S10**). When the HaRxL106 effector was expressed constitutively in the *bim1* mutant background, the resulting lines #1 and #3 were similar to *bim1* regarding petiole length, petiole insertion angle and chlorophyll content (**Figure 4B**, bar graphs). Furthermore, we examined the petiole length of three-week-old *bim123*/35S:YFP-HaRxL106 plants (**Figure 4C**). According to Yin *et al.* (2005), *bim123* plants were dwarf. This experiment revealed that this dwarfism is maintained even when expressing 35S:YFP-HaRxL106 in this background. Together, our results suggest that BIM1 is necessary for HaRxL106 to stimulate hypocotyl and petiole growth, as well as to increase leaf insertion angle. These HaRXL106-induced BIM1-dependent developmental modifications likely cause the observed constitutive SAS-like phenotype.

### HaRxL106 alters BIM1 activity generating an increase in the expression of BR-responsive genes

Collectively, our results pointed to a role of BIM1 as a susceptibility factor. One possible scenario was that the altered defense/immunity in *bim1* mutants is due to the reported function of BIM1 as a regulator of BR signaling (Liang *et al.* 2018). Thus, we wondered whether activation of this hormonal pathway occurred upon Hpa infection on susceptible wild type plants. We assessed the expression of known BR-responsive and SAS-related genes (*BR6OX, DWF4, ARF6, XTH19, IAA19*) (Sun *et al.*, 2010; Yu *et al.*, 2011; Bai *et al.* 2012; Oh *et al.* 2014; Liang *et al.*, 2018; Xu *et al.*, 2020) by qRT-PCR in wild type Col-0 seedlings infected with Hpa NoCo2. As shown in **Figure 5A**, at 96 hours after infection, the BR biosynthesis genes *BR6OX* and *DWF4*, as well as a gene linked to SAS responses mediated by BRs and auxins (*IAA19)* (Romanowski *et al.* 2021), were markedly induced in response to Hpa invasion.

**Figure 5:**
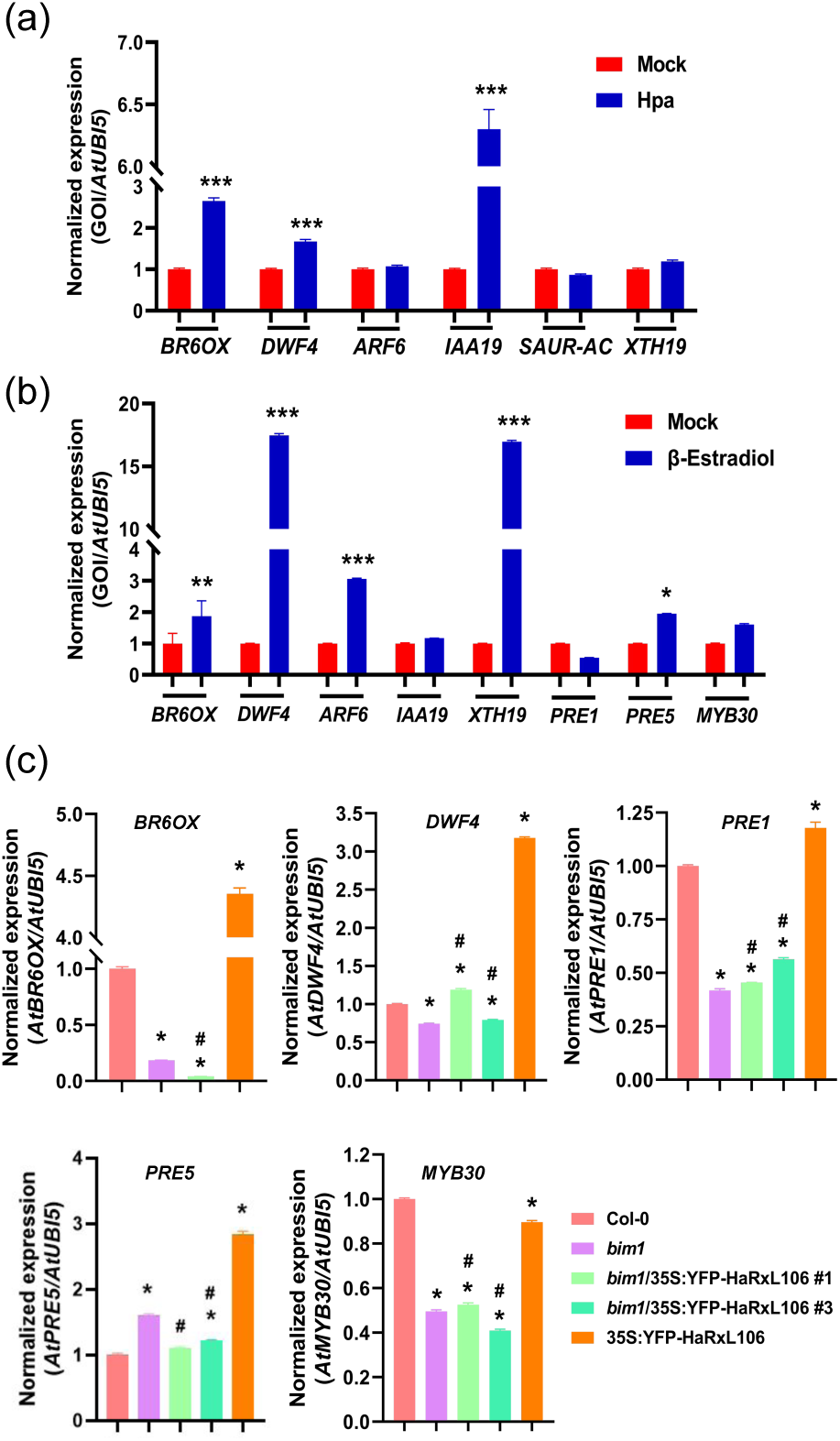
Hpa infection and HaRxL106 expression induce BR-responsive genes in a BIM1-dependent manner. **a.** Gene of interest (GOI) expression normalized to a reference gene (AtUBI5) at 96 hours post infection with Hpa NoCo2. Values are relative to each gene expression in uninfected plants of the same age (11-days-old). **b**. Normalized GOI expression in pER8:HA-HaRxL106 seedlings treated with DMSO (mock) or DMSO solution containing 1.25 μM β-Estradiol. Asterisks indicate statistically significant differences between treatments with *: p<0.05; **: p <0.01; ***: P<0.001 according to an ANOVA (Tukey post hoc test). **c**. Expression of the indicated genes normalized to the reference gene in 10 day-old seedlings of different genotypes. * indicates statistically significant differences regarding genotype Col-0; # indicates differences between *bim1*/35S:HaRxL106 plants and 35S:YFP-HaRXL106, according to a one-factor ANOVA (Tukey post hoc test, p<0.05).

Taking into account the above results, we then investigated if HaRxL106 was one of the effectors that Hpa deploys to alter the expression of the aforementioned genes. To avoid the known transcriptional feedback loop inhibition that controls the expression of many BR-responsive genes (Belkhadir *et al.* 2012), we decided to first study transcriptional changes of BR-responsive genes in Arabidopsis lines that expressed HaRxL106 under an estradiol-inducible promoter (Zuo *et al.*, 2000), we called these lines pER8:HA-HaRxL106 (**Supplementary Figure S11**). We observed that the induction of HaRxL106 expression stimulated the transcription of BR-responsive developmental genes as *BR6OX* and *DWF4*, as well as of *ARF6* and *XTH19,* genes that also respond to BRs-induced Auxins upregulation (Oh *et al.*, 2014; Xu *et al.* 2020) (**Figure 5B**). Interestingly, the expression pattern of BR-responsive genes was different from the one observed with Hpa infection (**Figure 5A**). This could be a consequence of the presence of other Hpa-delivered effectors or activation of PTI that possibly influence the expression of these genes.

Additionally, to ascertain if the expression of HaRxL106 could alter BIM1 activity as transcription factor, we also measured the expression of its direct targets, the transcription factors *PACLOBUTRAZOL RESISTANCE (PREs*) *PRE1, PRE5*, and *MYB30* (Yin *et al.*, 2002; Li *et al.*, 2009; Bai *et al.*, 2012; Liang *et al.* 2018; Buti *et al.* 2020). As it can be observed in **Figure 5B**, the effector induction affected the expression of BIM1 targets *PRE1*, *PRE5* and *MYB30* although to a lower extent, probably due to the tight positive and negative transcriptional regulation that these TFs are subject to (Zhang *et al.*, 2009).

Moreover, to determine if BIM1 was necessary for the activation of BR-responsive genes by HaRxL106, the expression levels of these genes were compared between Col-0 wild type seedlings, *bim1* mutants, and plants over-expressing HaRxL106 either in the wild-type or *bim1* mutant background. As shown in **Figure 5C**, constitutive expression of HaRxL106 in wild-type background induces the expression of *BR6OX, DWF4, PRE1* and *PRE5* (orange bars). On the contrary, seedlings that overexpress HaRxL106 in *bim1* background (green bars) exhibit levels of expression comparable to the *bim1* mutants (purple bars). In order to complement these data, we investigated how the binding of HaRxL106 to BIM1 might affect its direct activity as a transcription factor. For this, we performed promoter transactivation assays with a dual luciferase system that consists of a vector where a given promoter can be cloned to direct the expression of the firefly luciferase gene (Promoter X:fLUC). Following the previous sequence there is an in-frame construct to constitutively express the Renilla luciferase reporter (35S:rLUC). As reporters, we used the BIM1-specific promoters PRE5pro and SAUR-ACpro fused to the firefly luciferase gene (PRE5pro:fLUC and SAUR-ACpro:fLUC) built by Liang *et al.* (2018). BIM1-FLAG binding to the promoters of PRE5 and SAUR-AC genes has been previously reported by ChIP-qPCR (Liang *et al.*, 2018). We also collected data indicating that BIM1-FLAG upregulated the expression of SAUR-AC, while YFP-HaRxL106 slightly decreased SAUR-AC mRNA levels (**Supplementary Figure S12**). We then transiently expressed our promoter-reporter genes in *N. benthamiana* leaves via *A. tumefaciens*, co-infiltrating with or without the appropriate vectors to express BIM1-GFP and/or GFP-HaRxL106 and recorded the luminescence levels with a luminometer. Firefly luciferase activity (luminescence) was normalized to the Renilla Luciferase luminescence (fLUC/rLUC). As it can be observed in **Figure 6 A**, the PRE5 promoter is induced by BIM1-GFP and is also activated by GFP-HaRxL106, presumably acting through BIM1’s ortholog in *N. benthamiana*, but not by GFP alone. When both proteins, BIM1 and HaRxL106 are present, their effect is additive. Conversely, in the case of SAUR-AC (**Figure 6 B**) the negative effect of GFP-HaRxL106 is compensated by BIM1-GFP co-expression, while BIM1-GFP alone enhances the expression of SAUR-AC, in concordance with the data obtained by qRT-PCR (**Supplementary Figure S12)**. In summary, our results indicate that the transcription factor activity of BIM1 can be affected by HaRxL106 to effectively modulate a sector of the BR signaling pathway linked to the development of the SAS phenotype by up or downregulating different BR responsive genes.

**Figure 6:**
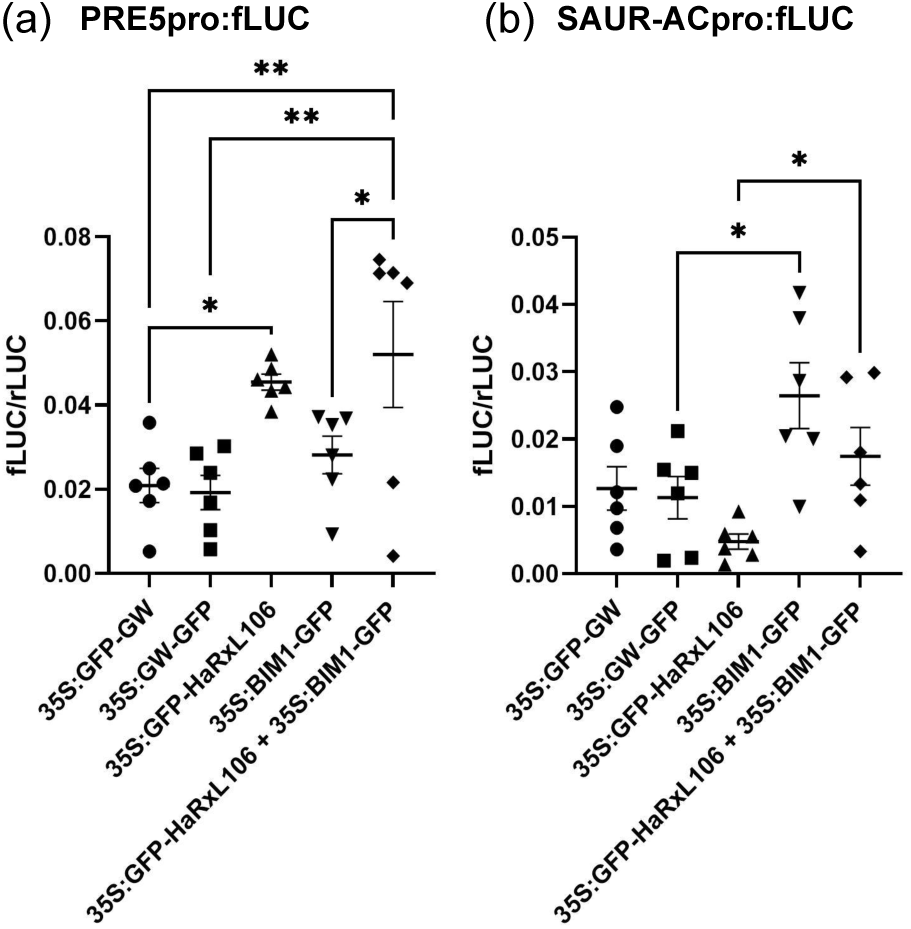
HaRxL106 affects the promoter activity of BIM1 targets. Transactivation assay in *N. benthamiana* leaves. *A. tumefaciens* AGL1 carrying the reporter plasmids (PRE5pro:LUC in **a** and SAUR-ACpro:LUC in **b.**) were co-infiltrated with AGL1 expressing the indicated constructs at OD=0.2 of each strain. 35S:GFP-GW and 35S:GW-GFP are the empty vectors controls where HaRxL106 and BIM1 CDS were cloned, respectively. Leaf tissues were harvested at 72 hpi and Firefly luciferase/Renilla luciferase ratio (fLUC/rLUC) was measured. Asterisks indicate statistically significant differences between treatments with *: p<0.05 and **: p <0.01 according to a parametric ANOVA (uncorrected Fisher’s LSD post hoc test) for a. and non-parametric ANOVA (uncorrected Dunn’s post hoc test) for b, n=6 for both panels.

## DISCUSSION

Cytoplasmic RxLR effector proteins interact with different types of plant targets (Fabro, 2022, Tör *et al.*, 2023). Many of the targets described to date are proteins directly or indirectly linked to plant immune responses and several are indeed manipulated by other types of effectors produced by bacterial or fungal pathogens (Mukthar *et al.*, 2011, Weßling *et al.* 2014, McLellan *et al.*, 2022). On the other hand, effector targets that participate in host metabolic processes other than immune responses have been less studied (Toruño *et al.* 2016). In this work we provide evidence that BIM1, a transcription factor involved in BR signaling, is targeted by the effector HaRxL106. We consider BIM1 as a plant susceptibility factor that HaRxL106 exploits to selectively induce the transcriptional activation of the BR signaling network. BR-dependent changes in gene expression correlate with morphological and physiological changes in aerial plant organs that generate a SAS-like phenotype. Plants with this phenotype show enhanced susceptibility not only to the obligate biotroph Hpa but also to a hemi-biotrophic bacteria (Pst), suggesting that the transcriptional alterations caused by HaRxL106 contribute to modify the morphology and probably the metabolic status of plant tissues making them more susceptible to different types of pathogens.

An analysis of our results in the context of the existing bibliography about effector targets roles, first led us to consider that BIM1 was a negative regulator of immunity, manipulated by HaRxL106 to suppress plant defense signaling. Previous RNAseq data of adult plants expressing HaRxL106 (HaRxL106-OE) infected with Pst revealed a suppression of SA-signaling (Wirthmueller *et al.* 2018), and the fact that *bim1* mutants were more resistant to Hpa and Pst fitted in this context. Other studies indicated that there is a negative crosstalk between the upregulation of BR-responsive transcription factors and PTI (Lozano-Durán *et al.* 2013, Malinovsky *et al.* 2014). Constitutive activation of BZR1 suppresses PAMP-triggered ROS production, PAMP-triggered gene expression and flg22-induced seedling growth inhibition. This suppression of PTI by BZR1 was particularly pronounced during fast seedling growth. Analysis of RNA-seq and ChIP data led those authors to suggest that BZR1 may interact with WRKY transcription factors that suppress PAMP-triggered ROS and downregulate transcription of defense genes. Furthermore, another BZR1 target gene, the bHLH transcription factor HBI1, negatively regulates the expression of PTI-marker genes (Fan *et al.* 2014; Malinovsky *et al.* 2014). In this sense, BIM1 functioning as a negative regulator of defenses might be mediated by one of its direct targets, the PRE TFs. It has been reported that PRE1 interacts with IBH1 (ILI1 BINDING BHLH 1). This protein inhibits HBI1. Upregulation of PRE1 would inactivate IBH1, thus increasing the activity of HBI1, causing a reduction in defense responses (Paik *et al.* 2017). However, we found that *bim1* mutants do not show an increased basal or infection-induced expression of SA-marker genes compared to the one observed in wild type and HaRxL106-OE plants (**Supplementary Figure S9**). Indeed, the phenotype of enhanced resistance we observed in *bim1* mutant plants (**Figure 2, Supplementary Figure S7**) occurs with a diminished level of PTI responses (flg22-triggered ROS and callose deposition **Figure 3**). Thus, it seems unlikely that BIM1 functions as a negative regulator of early PTI responses or defense gene transcription.

Another possibility was that HaRxL106-mediated BIM1-dependent transcriptional activation of certain BR-responsive genes (*BR6OX*, *DWF4*, **Figure 5**) would induce the synthesis of BR hormones, initiating and/or sustaining the execution of specific developmental programs like the SAS, deviating resources from defenses to growth and altering metabolic and physiological processes in the plant that may facilitate colonization. Plants that develop SAS decrease SA-dependent defenses, although the exact molecular mechanism is still unknown (de Wit *et al.* 2013). It is possible that actively dividing and elongating tissues (i.e. hypocotyls) display softer remodeled cell walls, easier to penetrate to establish haustoria, as well as perturbed nutrient pools that pathogens may thrive on. Indeed, the increase in chlorophyll content that we observe in HaRxL106-OE plants (**Figure 4B**) could be indicating enhanced photosynthesis as it has been reported to occur surrounding infection sites of the oomycete *Albugo candida* in Arabidopsis at early stages of interaction (Chou *et al.* 2000). Enhanced photosynthesis produces changes in soluble carbohydrates availability that can be used for pathogen nutrition and growth (Berger *et al.* 2007, Herlihy *et al.*, 2019). In support of this hypothesis, plants that overexpress BIM1-FLAG show some of the early developmental features of HaRxL106-OEs (e.g. petiole length; **Figure 4C**) and also exhibit enhanced susceptibility to Hpa (**Figure 2A, Supplementary Figure S7B**). Additionally, it was previously observed that endogenous induction of BR signaling by overexpressing the receptor BRI1 (BRI1-OE), or its hypermorphic allele BRI1^sud1^, promotes the SAS-like growth and increased susceptibility to Hpa virulent race NoCo2. This enhanced sporulation is more pronounced on the first true leaves, organs in which BR-driven cell elongation programs are hyperactive (Belkhadir *et al.* 2012). Arabidopsis BRI1-OE, mutants BRI1^sud1^ or those constitutively expressing *DWF4*, a gene encoding the BR biosynthetic enzyme C-22 hydroxylase, display elongated petioles and narrow curved leaf laminas and show reduced flg22-induced ROS burst and callose deposition (Belkhadir *et al.* 2012). Interestingly, these phenotypes are all present in HaRxL106-OE plants.

The detailed molecular mechanism by which HaRxL106 promotes the transcriptional activity of BIM1 still remains elusive and merits further investigation. BIM1 is post translationally modified by phosphorylation (PhosPhAt, https://www.psb.ugent.be/webtools/ptm-viewer/protein.php?id=AT5G08130.5 and sumoylation (Miller *et al.* 2010). These protein modifications might be modulated by HaRxL106 affecting BIM1 function as a transcription factor. Alternatively, the effector might be altering BIM1 subcellular localization, retaining it into the nucleus, or stabilizing BIM1 binding to the DNA or modulating BIM1-BES1 heterodimer formation or even acting as a scaffold for the recruitment of co-transcriptional regulators, like other HaRxL106-interactors, such as RCD1 (Wirthmueller *et al.* 2018). RCD1 localizes to the plant cell nucleus and binds several transcription factors (Jaspers *et al.* 2009). Loss of RCD1 affects plant development and several stresses signaling pathways causing alterations in plant growth and development, likely mediated by its interaction with PHYTOCHROME INTERACTING FACTORS (PIFs) 3, 5 and 7 (Jaspers *et al.*, 2009). In fact, HaRxL106-mediated SAS-like phenotype is also largely suppressed in *rcd1*, and both *bim1* and *rcd1* mutants are impaired in activating the SAS developmental program (Cifuentes Esquivel *et al.* 2013; Salazar, 2010). It remains possible that HaRxL106 interacts with both targets at the same time in the plant host nucleus to promote alterations in plant development (via BIM1) while causing SA-pathway dependent defense suppression (via RCD1) (see graphical abstract model). Whether these interactions are mutually exclusive and whether they act additively or synergistically to promote host colonization deserves further investigation. In our hands, Y2H assays indicate that both effector targets, RCD1 and BIM1, can indeed interact in the absence of HaRxL106 (**Supplementary Figure 13**). Future experiments will help us to determine if this interaction can be altered by the presence of HaRxL106.

Targeting of transcription factors (TFs) involved in plant growth and development has been previously reported for oomycete effectors. Hpa effector HaRxL21 expression in Arabidopsis or *N. benthamiana* generates enhanced susceptibility to Hpa or *P. infestans*, respectively (McLellan *et al.* 2022). This effector interacts with the transcriptional co-repressor TOPLESS (TPL) which participates in numerous processes of plant development. Arabidopsis lacking TPL show enhanced susceptibility to Pst, Hpa and the fungus *Botrytis cinerea* (Zhu *et al.* 2010, Harvey *et al.* 2020). Similarly, HaRxLL470, interacts with ELONGATED HYPOCOTYL 5 (AtHY5) a TF involved in photomorphogenesis. The effector reduces HY5 binding to DNA, compromising transcription of defense marker genes (Chen *et al.* 2021). *P. infestans* effector Pi03192 and *Bremia lactucae* effectors BRLs 04,05,08 and 09 target NAC (NAM/ATAF/CUC) TFs involved in germination and stress responses, preventing their translocation to the nucleus and hindering host defenses (McLellan *et al.* 2013; Meisrimler *et al.* 2019). The *P. capsici* effector CRN12-997 binds a tomato TCP (Teosinte-Branched-Cycloidea and PCF 14, SlTCP14-2), inhibiting its binding to DNA and immunity-associated activity (Stam *et al.* 2021). The ortholog of TCP14 in Arabidopsis positively regulates photo-and thermo-morphogenesis, and is targeted by Hpa effector HaRxL45, but the consequences of this interaction have not been described (Yang *et al.* 2017). Regarding manipulation of TFs specifically involved in the BR-pathway, there is only one previous report, the effector AVR2 of *Phytophthora infestans,* which upregulates potato TF StCHL1, a positive regulator of BR signaling. Potato plants over-expressing AVR2 show an altered growth morphology, increased BR marker gene expression and enhanced susceptibility to *P. infestans* (Turnbull *et al.* 2017). In all the above-mentioned examples, the consequences that effector-target interactions had over different types of plant defense responses were clearly established, supporting the hypothesis that the crosstalk existing between growth and defenses is skillfully exploited by adapted pathogens. On the contrary, the influences that effector-target interaction had on plant growth/development were not investigated. We consider that our studies can contribute to clarify how effectors are able to manipulate the developmental pattern of the plant host and the consequences this has on pathogen fitness.

Regarding the proposed role of BIM1 as a susceptibility factor and envisaging possible application of these findings to plant disease management, it remains to be determined if economically relevant pathogens also require BIM1 to establish a compatible interaction. BIM1 is conserved across the green plant lineage (*Viridiplantae*) (https://phylogenes.arabidopsis.org/tree/PTHR46412). Genes that encode BIM1 homologs/orthologs are found in crop plants such as cotton, cocoa, castor bean and walnut. In tomato, the closest sequence to BIM1, *SlBIM1a*, has been proposed as a negative regulator of cell size in fruit pericarp cells. Indeed, *SlBIM1a* silencing specifically increased fruit and pericarp cell size, as well as plant stem length (Mori *et al.*, 2021). Other orthologs are found in soybean, wheat, sorghum, sunflower, rapeseed, grapevine and other edible species whose economic value does not lie in the flower/fruit/seed such as cabbage, lettuce, spinach and potato (*Solanum tuberosum*). In this last species, there is a gene annotated as transcription factor *StBIM1* (ID:PGSC0003DMG400024523; The Potato Genome Sequencing Consortium, 2011). Interestingly, it was recently reported that transient expression of HaRxL106 in *N. benthamiana* leaves promoted susceptibility to *P. infestans,* an oomycete of huge economical relevance (McLellan *et al.* 2022). Thus, it might be worth studying the role of BIM1 in the *P. infestans*-potato or tomato interaction. Additionally, to reliably determine if the elimination or modification of BIM1 does not have deleterious effects in plant growth and yield, it is necessary to carry out analyzes that investigate BIM1 involvement in flowering and seed production. Absence of BIM1 in Arabidopsis does not cause major growth pattern alterations in plant shape or size (**Figure 4**) but we observed an early flowering phenotype. This is consistent with a described function of BIM1 on reproductive development, where together with BES1, it binds to a BRRE *cis*-element in the first intron of the key floral repressor FLOWERING LOCUS C (FLC). BES1/BIM1 binding recruits a histone demethylase to promote FLC expression, blocking floral transition and favoring vegetative development (Li *et al.* 2018, Li and He, 2020). Xing *et al.* (2013) described that *bim1* mutants are only slightly less fertile than wild Col-0 plants. Likewise, *bim1* mutants show minor defects in embryo development (Chandler *et al.* 2009). If it turns out that BIM1 orthologs in *Solanaceae* are required for susceptibility to oomycetes or other pathogens, then editing BIM1 in potato, which reproduces clonally through its tubers, constitutes an attractive strategy to achieve greater resistance without the energetic cost associated with implementing active defense responses (Wang *et al.*, 2007; Canet *et al.* 2010; Groszmann *et al.* 2015; van Butselaar and Van den Ackerveken, 2020). We consider that our results could be a starting point for potential biotechnological applications in order to improve cultivar defenses without dampening growth, thus supporting food security.

## EXPERIMENTAL PROCEDURES

### Plant material and growth conditions

Wild type Arabidopsis seeds ecotype Columbia-0 (Col-0) wild type and 35S:YFP-HaRxL106 were described previously (Fabro *et al.*, 2011; Wirthmueller *et al.* 2015, 2018). In the case of *bim1* mutants, we first obtained seeds of allegedly *bim1* mutant plants from Yin *et al.* 2005 and also from the ABRC (line SALK_085924). After careful genotyping we found out that this line did not carry the expected T-DNA insertion in the BIM1 ORF, while another line requested to Dr. John Chandler, SALK_132178, was a homozygous mutant, and thus in this work we refer to this line as *bim1* mutant (**Supplementary Figure S6A, B, C**). We also quantified BIM1 expression in this mutant line by RT-qPCR (**Supplementary Figure S6D**). *bim1* mutants expressed a transcript to a significantly lower level (p<0.001) than wild plants. Even when the expression of BIM1 in the mutants is not zero; the transcript detected is likely to be truncated as the primers used for the PCR hybridize upstream of the T-DNA inserted at the end of the 9^th^ exon. The *bim123* triple mutant from Yin *et al.*, 2005 was developed with SALK_085924 line (insertion in BIM1 CDS at 5’ end, 1^st^ exon) crossed by SALK_074689C: insertion in BIM2 and SALK_079683: insertion in BIM3. The BIM1-FLAG overexpressing line was obtained from Dr. Hongtao Liu (Liang *et al.*, 2018). *bim23* double mutants were obtained by genotypic analysis of presumed *bim123* triple mutants sent by the ABRC, deposited there by the authors of Yin *et al.*, 2005. It turned out that many of these plants had the wild type allele of BIM1 gene. We selected homozygous mutants for the insertion on the BIM1 5’ end using primers designed to screen the SALK_085924 line (**Supplementary Figure S6E**). We called these lines *bim123* and are the ones we complemented with BIM1-FLAG or with HaRxL106. A summary of these data is presented in Supplementary Table 1.

Stable Arabidopsis 35S:GFP-HaRxL106 expressing lines were developed as follows: pENTR4 clone containing HaRxL106 CDS (Wirthmueller *et al.* 2018) was recombined in pK7WGF2 destination vector (Ghent University) by Gateway cloning using LR clonase II (Invitrogen), obtaining GFP-HaRxL106 expression clones resistant to spectinomycin. Transgenic plants were developed according to the floral dip method (Clough and Bent, 1998). Col-0 wild type plants were dipped in a suspension containing 5% sucrose, 0.05% Silwet L77 and *Agrobacterium tumefaciens* GV3101 carrying these vectors. Transgenic lines were selected by their kanamycin resistance on MS Agar plates over 3 generations (T3s). Similarly, *bim1(23)*/35S:YFP-HaRxL106 lines were made by dipping the inflorescences of *bim1, bim23* or *bim123* plants in a suspension of *A. tumefaciens* carrying 35S:YFP-HaRxL106, and selected by their glufosinate/BASTA resistance.

For hypocotyl measurements and estradiol induction experiments, Arabidopsis seeds were sterilized in Triton X-100 0.05%, ethanol 70% for 10 minutes, then washed two times in ethanol 96% for 5 minutes and dried in a laminar flow chamber. They were sown in 0.5x Murashige-Skoog medium (MS), 1% agar plates, pH=5.7, and stratified at 4°C for 2-5 days. Plates were placed vertically in a growth chamber with 12:12 h photoperiod, 22°C and 70% relative humidity and 5500 lux light intensity (LED warm light, 6500 K), and hypocotyl length was measured after 7 days using ImageJ software. For estradiol treatment, a 5 mM stock solution was prepared in DMSO, which was diluted in MQ water to a final concentration of 1.25 µM. One ml of this was used to spray the seedlings in each 90 mm MS plate. This treatment was applied 5 times every 48 hours, from day 2 after germination. On day 11, seedlings were harvested, total RNA was extracted and 2 μg used for reverse transcription and gene expression assessed by qPCR as described below.

For other experiments, seeds were stratified for 2-5 days at 4°C, directly sown in a commercial substrate (Grow Mix Multipro, Terrafertil SA) and placed in a plant growth chamber (Demetra 380L, J3 Desarrollos, Mar del Plata, Argentina) with 12:12 photoperiod, 22°C (except for Hpa infection experiments in which plants were grown at 18°C), 70% humidity and 7000 lux. The same growth conditions were used for *N. benthamiana* plants. Images of 2-3-week-old plants were used to measure petiole length by ImageJ. In 8-week-old adult plants, angles of insertion to the rosette were measured using a protractor, then the leaves were detached and the petioles measured with a caliper.

### RNA purification and qPCR

RNA was purified using the SDS-LiCl RNA method (Verwoerd *et al.* 1989) and reverse transcription performed as previously described (Cambiagno *et al.* 2015). TransStart® Green qPCR SuperMix (Transbiotec), was used to perform qPCR with a first denaturation step at 95°C for 3 minutes, then 40-50 cycles of 95°C for 20 sec, annealing at 58°C for 30 sec and extension at 72°C for 15 sec. *AtUBI5* (Ubiquitin 5; *At3g62250*) was used as a reference gene. Specific primers for defense and BR-related marker genes are listed in Supplementary table 2. The amplification efficiency for each well was calculated using LinRegPCR Software (Ramakers *et al.*, 2003), then the average efficiency (E) for each primer pair was used to estimate the relative quantity according to: E^(Ct^ ^control^ ^sample-Ct^ ^test)^. The normalized expression of the genes of interest (GOI) is the relative quantity of the GOI/relative quantity of the reference.

### Constructs used for BiFC,Co-IP and transactivation assays

The constructs of HaRxL106 full length, SV40NLS-HaRxL106-ΔC and HaRxL106Cterm58 cloned in pENTR-D-TOPO were generated previously (Wirthmueller *et al.*, 2015) (Supplementary Table 1). In this work we used those pENTR clones to recombine in different destination vectors. For BiFC, Gateway-compatible pAM-PAT-35S-YFPN/C plasmids were used (Lefebvre *et al.* 2010), leaving the tag (YFPn) at the N-terminal domain of the effector and at the C-terminal (YFPc) at the C-term of the interactor, in order not to interfere with the functionality and subcellular localization of these proteins. BIM1-CFPc and BIM1-FLAG plasmids were kindly sent by Dr. Hongtao Liu (Liang *et al.* 2018). BIM2 and BIM3 CDS were cloned into pENTR4 by Gibson cloning using the primers listed on Supplementary Table 2. These pENTR4 clones were used to recombine into pAM-PAT-35S-YFPc with the fluorescent tag at the C-term of the protein. For CoIP, we tagged the different versions of HaRxL106 at its N-terminal with GFP using the destination vector pK7WGF2. For the transactivation assyt, BIM1 was cloned into pENTR-D-TOPO amplifying the full-length sequence from the BIM1-CFPc plasmid, and then recombined with pB7FWG2. Destination vectors were introduced in *E. coli* for amplification, purification, sequence verification and then transformed into *A. tumefaciens* GV3101 to develop transient and stable transformation of plants.

### Transient expression of proteins *in planta*

*N. benthamiana* plants (1-month-old) were infiltrated with a suspension of 10 mM MgCl_2_, 10 mM MES buffer pH=5.7, 150 μM acetosyringone and the *A. tumefaciens* carrying the above-mentioned plasmids. When two proteins were co-expressed, the suspension contained OD_600_=0.3 of each strain; if they were expressed alone the OD_600_ was 0.4. In all cases, *A. tumefaciens* carrying P19 silencing suppressor was co-infiltrated at OD_600_ 0.05. Confocal microscopy images were taken after 48-72 hours post infiltration.

### CoIP assays

Total protein contents were isolated using extraction buffer (50 mM Tris-HCl pH=7.4, 150 mM NaCl, 10% (v/v) glycerol, 5 mM DTT, 1× plant PIC, 0.2% NP-40, 10 μM MG132, 100 μg/ml DNAse, 5 mM CaCl_2_ and 10 mM MgCl_2_). GFP-tagged proteins were immunoprecipitated by an *ad hoc* produced anti-GFP nanobody (pOPINE GFP nanobody:Halo:His6; Addgene plasmid #111090; Chen *et al.* 2018) covalently bound to magnetic beads (Magne HaloTag® Beads, Promega). Total extracts and IPs were run in SDS-PAGE and western blot was performed using anti GFP (Polyclonal, made in rabbit, #3999-100, AMSBIO, AMS biotechnology -Europe-Ltd) and anti-FLAG (ANTI-FLAG® M2, monoclonal mouse, #F3165-2MG, Sigma Aldrich) antibodies. Secondary antibodies were LICOR IRDye® 800CW anti-rabbit IgG -926-32211-or anti-mouse IgG-926-32210-and were used at 1:10.000 dilution. WB membranes were scanned with an Odyssey Infrared Imaging System (LI-COR, Inc., Lincoln, NE, USA).

### Transactivation assay

Dual luciferase assay was performed in. *N. benthamiana* leaves. *A. tume*f*aciens* AGL1 carrying the reporter plasmids (PRE5pro:LUC and SAUR-ACpro:LUC, Liang *et al.* 2018) were co-infiltrated with AGL1 expressing the empty vectors 35S:GFP-GW (pK7WGF2) and 35S:GW-GFP (pB7FWG2) or the 35S:GFP-HaRxL106 and 35S:BIM1-GFP constructs at OD=0.2 each in 10 mM MgCl_2_, 10 mM MES buffer pH=5.7, 150 μM acetosyringone. Three days later (72 hpi), 6 leaf discs (0.23 cm^2^) of each combination were flash frozen in liquid nitrogen and treated subsequently as technical replicates. Tissues were ground and the Firefly luciferase/Renilla luciferase ratio (fLUC/rLUC) assessed using the Dual-Luciferase® Reporter Assay System (Promega) according to manufacturer instructions.

### Yeast Two Hybrid experiments

The Y2H assay was carried out using the ProQuest^TM^ Two-Hybrid System (Invitrogen). For this purpose, pENTR4 containing the full length BIM1 CDS (1599 bp, splicing variant 5, At5g08130.5) or the N-terminal domain of BIM1 (835 bp) were recombined via Gateway in the pDEST22 prey vector. To clone into the pDEST32 bait vector we used full-length construct of HaRxL106 as well as the N-terminal domain (SV40NLS-HaRxL106ΔC) or C-terminal domain (HaRxL106-c58) (Wirthmueller *et al.*, 2015 and 2018). These plasmids were introduced into the yeast strain Ma203V according to the manufacturer instructions, and the transformed cells selected in a -WL commercial medium (SC-WL, Synthetic Defined Yeast Media, MP Biomedicals™). Subsequently, they were plated on the selective medium for clones carrying both plasmids (SC-WL) and media to select clones expressing interactive proteins (SC-WLH) containing different concentrations of 3-Aminotriazole. Ten and 25 mM 3-AT were used for all experiments but sometimes only one concentration is shown in figures. The plates were incubated at 28-30 °C and photographed after 2-3 days. As positive controls, we used Krev1 and RalGDS following manufacturer instructions (Invitrogen, USA). As negative control mutant (^m^) versions of Krev1 and RaIGDS that do not interact were used. Additional negative controls were generated to monitor autoactivation, by co-transforming yeasts with combinations of one interactor and one empty vector, or both empty vectors.

### In vitro Pull-Down assay

The full-length sequence of BIM1, cloned into pMAL5c to generate MBP-BIM1 was obtained from Dr. Hongtao Liu (Liang *et al.*, 2018). This recombinant protein was expressed in *E. coli* BL21 and affinity purified using amylose resin (NEB #E8021L) as previously described for other MBP tagged proteins (Liang *et al.*, 2018; Fabro *et al.* 2020). HIS-HaRxL106 cloned into pOPINF was expressed into SoluBL21 and purified as previously described (Wirthmueller *et al.*, 2015) using a Ni-NTA resin (Probond, Invitrogen 46-0019).The purified effector protein was applied to the column with the amylose bound MBP or MBP1-BIM1 and let interact for 1 hour in binding buffer (Tris-HCl 50 mM pH=7.4, 150 mM NaCl, 1 mM EDTA, 0.2 %Triton X-100, 10 % glycerol and 0.5 mM PMSF). Then four 10 X column volume washes were performed with the same binding buffer and finally proteins were eluted with 15 mM Maltose. A 30 μl fraction of the washed amylose beads and a concentrated eluate (10 X, performed via a10 KDa cut off-Amicon ultracel filter) were loaded onto two twin SDS-PAGE gels. One gel was stained with coomassie brilliant blue to reveal MBP and MBP-BIM1 and the other was transferred to a PVDF membrane perform western blot analysis using primary antibody anti-HIS made in mouse (SIGMA, H1029 dilution 1:3000) and the secondary antibody anti mouse IgG used for the CoIPs.

### Hpa and Pst infection and quantification

*Hyaloperonospora arabidopsidis* isolate NoCo2 conidiospores at a concentration of 1-5x10^5^ spores/ml were inoculated on 7/10-day-old Arabidopsis seedlings of different genotypes. The seedlings stayed 7-10 days in a high humidity plastic dome and conidiospore proliferation was counted using a Neubauer chamber and made relative to the collected seedlings’ fresh weight. For *Pseudomonas syringae* pv. *tomato* (DC3000, Pst) infection assays, 1.5 to 2-month-old plants were inoculated at a concentration of 5 x 10^5^ or 10^6^ CFU/ml in 10 mM MgCl_2_, infiltrating them through the abaxial side of the leaves with a needleless syringe. At 1 and 3 dpi, infected leaf discs were cut, ground in 10 mM MgCl_2_ and serial dilutions were plated in LB Agar plates supplemented with 100 μg/ml Rifampicin and 50 μg/ml Kanamycin. Bacterial titers were counted after 48-72 hours incubation at room temperature.

### Chlorophyll quantitation

To estimate the total chlorophyll content, leaf discs of 0.307 cm^2^ (cork borer #2) were cut, discolored on 70% ethanol at 80 °C for 10 minutes and centrifuged at 14,000 rpm for 5 minutes. The optical density of the supernatants was read at 654 nm following the protocol proposed by Vernon (1960).

### Statistical analysis

Statistical analyses were performed using InfoStat (Balzarini *et al.* 2008) and Graph Pad Softwares (GraphPad Software Inc.; San Diego, CA, USA).

## ACCESSION NUMBERS

Please see Supplementary table 1.

## ACKNOWLEDGEMENTS

To Dr. Hongtao Liu and Dr. John Chandler for *bim1* and *bim123* mutant seeds. To Dr. Hongtao Liu for BIM1-FLAG seeds and several constructs: BIM1-CFPc/FLAG,MBP-BIM1 and PRE5/SAUR-ACpro:LUC. To Dr. Dae Sung Kim, from Dr. Jonathan D.G. Jones laboratory (TSL, Norwich, UK) for sharing unpublished data on HaRxL106 interactors. To Prof. Dr. Tina Romeis for supporting the research stays of G.F. and M.F.B. at Free University of Berlin and IPB Halle. To Dr. Maria Elena Alvarez for the support to G. F and the initial PhD fellowship of M.F.B. To Dr. Nicolas Cecchini for his critical and comprehensive reading of the manuscript. To Dr. Maria Florencia Nota, Dr. Ignacio Lescano, Dr. Jessica Erickson, Dr. Elvio Benatto, Susanne Kirsten and Sylvia Krueger for their technical and experimental assistance. To Drs. Cecilia Sampedro and Carlos Mas from CEMINCO for their assistance with confocal microscopy. To undergrad student Sol Angulo for her assistance in sample preparation. The authors declare no conflicts of interest.

## REFERENCES

Albrecht, C., Boutrot, F., Segonzac, C., Schwessinger, B., Gimenez-Ibanez, S., Chinchilla, D., Rathjen, J. P., De Vries, S. C., & Zipfel, C. (2012). Brassinosteroids inhibit pathogen-associated molecular pattern-triggered immune signaling independent of the receptor kinase BAK1. Proc. Natl Acad. Sci. USA, 109, 303–308. 10.1073/pnas.1109921108

Bai, M. Y., Shang, J. X., Oh, E., Fan, M., Bai, Y., Zentella, R., Sun, T. P., and Wang, Z. Y. (2012). Brassinosteroid, gibberellin and phytochrome impinge on a common transcription module in Arabidopsis. Nature Cell Biology. 14, 810–817. 10.1038/ncb2546.

Balzarini, M., Gonzalez, L., Tablada, E., Casanoves, F., Di Rienzo, J., and Robledo, C. (2008). Manual del Usuario InfoStat Software Estadístico. Infostat, 53(January), 336.

Belkhadir, Y., Jaillais, Y., Epple, P., Balsemão-Pires, E., Dangl, J. L., & Chory, J. (2012). Brassinosteroids modulate the efficiency of plant immune responses to microbe-associated molecular patterns. Proc. Natl Acad. Sci. USA 109, 297–302. 10.1073/pnas.1112840108

Belkhadir Y. and Jaillais Y.(2015) The molecular circuitry of brassinosteroid signaling. New Phytol. 206, 522–40. doi: 10.1111/nph.13269.

Belkhadir Y, Wang X, Chory J. (2006) Arabidopsis brassinosteroid signaling pathway. Sci STKE. 364, cm5. doi: 10.1126/stke.3642006cm5.

Berger S., Sinha A.K. and Roitsch T. (2007) Plant physiology meets phytopathology: plant primary metabolism and plant-pathogen interactions. J Exp Bot. 58, 4019–26. doi: 10.1093/jxb/erm298.

Boevink PC, Birch PRJ, Turnbull D, Whisson SC. (2020) Devastating intimacy: the cell biology of plant-Phytophthora interactions. New Phytol. 228, 445–458. doi: 10.1111/nph.16650.

Bücherl, C. A., Jarsch, I. K., Schudoma, C., Segonzac, C., Mbengue, M., Robatzek, S., MacLean, D., Ott, T., and Zipfel, C. (2017) Plant immune and growth receptors share common signalling components but localise to distinct plasma membrane nanodomains. Elife 6, e25114

Buti, S., Hayes, S. and Pierik, R. (2020). The bHLH network underlying plant shade-avoidance. Physiologia Plantarum. 169, 312–324. 10.1111/ppl.1307410.1111/ppl.13074

Caillaud, M.C., Wirthmueller L, Fabro G, Piquerez SJ, Asai S, Ishaque N, Jones JD. (2012) Mechanisms of nuclear suppression of host immunity by effectors from the Arabidopsis downy mildew pathogen Hyaloperonospora arabidopsidis (Hpa). Cold Spring Harb Symp Quant Biol.77:285–93. doi: 10.1101/sqb.2012.77.015115.

Cambiagno, D. A., Lonez, C., Ruysschaert, J. M., & Alvarez, M. E. (2015). The synthetic cationic lipid diC14 activates a sector of the A rabidopsis defence network requiring endogenous signalling components. Molecular plant pathology, 16(9), 963–972.

Canet, J. V., Dobón, A., Ibáñez, F., Perales, L., and Tornero, P. (2010). Resistance and biomass in Arabidopsis: A new model for Salicylic Acid perception. Plant Biotechnology Journal. 8, 126–141. 10.1111/j.1467-7652.2009.00468.x

Chandler, J. W., Cole, M., Flier, A., & Werr, W. (2009). BIM1, a bHLH protein involved in brassinosteroid signalling, controls Arabidopsis embryonic patterning via interaction with DORNRÖSCHEN and DORNRÖSCHEN-LIKE. Plant Molecular Biology. 69, 57–68. 10.1007/s11103-008-9405-6

Chen C., Masi R., Lintermann R. and Wirthmueller L.(2018) Nuclear Import of Arabidopsis Poly(ADP-Ribose) Polymerase 2 Is Mediated by Importin-α and a Nuclear Localization Sequence Located Between the Predicted SAP Domains. Front Plant Sci. 9, 1581. doi: 10.3389/fpls.2018.01581.

Chen S., Ma T., Song S., Li X., Fu P., Wu W., Liu J., Gao Y., Ye W., Dry I.B. and Lu J.(2021) Arabidopsis downy mildew effector HaRxLL470 suppresses plant immunity by attenuating the DNA-binding activity of bZIP transcription factor HY5. New Phytol. 230, 1562–1577. doi: 10.1111/nph.17280.

Chinchilla, D., Shan, L., He, P., de Vries, S., and Kemmerling, B. (2009) One for all: the receptor-associated kinase BAK1. Trends Plant Sci. 14, 535–541

Chou H.M., Bundock N., Rolfe S.A. and Scholes J.D. (2000) Infection of Arabidopsis thaliana leaves with Albugo candida (white blister rust) causes a reprogramming of host metabolism. Mol Plant Pathol. 1, 99–113. doi: 10.1046/j.1364-3703.2000.00013.x.

Cifuentes-Esquivel, N., Bou-Torrent, J., Galstyan, A., Gallemí, M., Sessa, G., Salla Martret, M., Roig-Villanova, I., Ruberti, I., and Martínez-García, J. F. (2013). The bHLH proteins BEE and BIM positively modulate the shade avoidance syndrome in Arabidopsis seedlings. Plant Journal, 75, 989–1002. 10.1111/tpj.12264

Clough S.J. and Bent A.F. (1998) Floral dip: a simplified method for Agrobacterium-mediated transformation of Arabidopsis thaliana. Plant Journal 16, 735–43. doi: 10.1046/j.1365-313x.1998.00343.x.

DeFalco T.A. and Zipfel C. (2021) Molecular mechanisms of early plant pattern-triggered immune signaling. Mol Cell. 81, 3449–3467. doi: 10.1016/j.molcel.2021.07.029.

De Wit, M., Spoel, S.H., Sanchez-Perez, G.F., Gommers, C.M.M., Pieterse, C.M.J., Voesenek, L.A.C.J., and Pierik, R. (2013). Perception of low red: far-red ratio compromises both salicylic acid-and jasmonic acid-dependent pathogen defences in Arabidopsis. Plant J. 75, 90–103,

Dodds, P.N. (2023) From Gene-for-Gene to Resistosomes: Flor’s Enduring Legacy Mol Plant Microbe Interact. 36, 461–467. doi: 10.1094/MPMI-06-23-0081-HH.

Dong, S., and Ma, W. (2021). How to win a tug-of-war: the adaptive evolution of Phytophthora effectors. Current Opinion in Plant Biology, 62, 102027. 10.1016/j.pbi.2021.102027

Fabro G., Steinbrenner J., Coates M., Ishaque N., Baxter L., Studholme D.J., Körner E., Allen R.L., Piquerez S.J., Rougon-Cardoso A., Greenshields D., Lei R., Badel J.L., Caillaud M.C., Sohn K.H., Van den Ackerveken G., Parker J.E., Beynon J. and Jones J.D. (2011) Multiple candidate effectors from the oomycete pathogen Hyaloperonospora arabidopsidis suppress host plant immunity. PLoS Pathog. 7, e1002348. doi: 10.1371/journal.ppat.1002348.

Fabro G. Oomycete intracellular effectors: specialised weapons targeting strategic plant processes. (2022) New Phytol. 233, 1074–1082. doi: 10.1111/nph.17828.

Fabro G, Cislaghi AP, Condat F, Deza Borau G, Alvarez ME. (2020) The N-terminal domain of Arabidopsis proline dehydrogenase affects enzymatic activity and protein oligomerization. Plant Physiol Biochem;154:268–276. doi: 10.1016/j.plaphy.2020.04.019

Fan M., Bai M.-Y., Kim J.-G., Wang T., Oh E., Chen L., Park C. H., Son S.-H., Kim S.-K., Mudgett M. B. et al. (2014). The bHLH transcription factor HBI1 mediates the trade-off between growth and pathogen-associated molecular pattern–triggered immunity in Arabidopsis. Plant Cell 26, 828–841. 10.1105/tpc.113.121111

Figueroa M., Ortiz D., Henningsen E.C. (2021) Tactics of host manipulation by intracellular effectors from plant pathogenic fungi. Curr Opin Plant Biol. 62,102054. doi: 10.1016/j.pbi.2021.102054.

Groszmann, M., Gonzalez-Bayon, R., Lyons, R. L., Greaves, I. K., Kazan, K., Peacock, W. J., & Dennis, E. S. (2015). Hormone-regulated defense and stress response networks contribute to heterosis in Arabidopsis F1 hybrids. Proc Natl Acad Sciences USA, 112, E6397–E6406. 10.1073/pnas.1519926112

Han, X., and Kahmann, R. (2019). Manipulation of phytohormone pathways by effectors of filamentous plant pathogens. Front in Plant Science. 10.3389/fpls.2019.00822

Harvey S., Kumari P., Lapin D., Griebel T., Hickman R., Guo W., Zhang R., Parker J.E., Beynon J., Denby K. and Steinbrenner J.(2020) Downy Mildew effector HaRxL21 interacts with the transcriptional repressor TOPLESS to promote pathogen susceptibility. PLoS Pathog. 16, e1008835. doi: 10.1371/journal.ppat.1008835.

He Q., McLellan H., Boevink P.C. and Birch P.R.J.(2020) All Roads Lead to Susceptibility: The Many Modes of Action of Fungal and Oomycete Intracellular Effectors. Plant Commun. 24, 100050. doi: 10.1016/j.xplc.2020.100050.

Herlihy J., Ludwig N.R., van den Ackerveken G. and McDowell J.M.. (2019) Oomycetes Used in Arabidopsis Research. Arabidopsis Book. 17:e0188. doi: 10.1199/tab.0188.

Huot B., Yao J., Montgomery B.L. and He S.Y.(2014) Growth-defense tradeoffs in plants: a balancing act to optimize fitness. Mol Plant. 7, 1267–1287. doi: 10.1093/mp/ssu049.

Igarashi D., Tsuda K. and Katagiri F. (2012)The peptide growth factor, phytosulfokine, attenuates pattern-triggered immunity. Plant J. 71, 194–204. doi: 10.1111/j.1365-313X.2012.04950.x.

Jaspers P., Blomster T., Brosché M., Salojärvi J., Ahlfors R., Vainonen J.P., Reddy R.A., Immink R., Angenent G., Turck F., Overmyer K. and Kangasjärvi J. (2009) Unequally redundant RCD1 and SRO1 mediate stress and developmental responses and interact with transcription factors. Plant Journal. 60, 268–79. doi: 10.1111/j.1365-313X.2009.03951.x.

Judelson H. S. and Ah-Fong A. M. V. (2019) Exchanges at the Plant-Oomycete Interface That Influence Disease, Plant Physiology,179, 1198–1211, 10.1104/pp.18.00979

Kang S., Yang F., Li L., Chen H., Chen S. and Zhang J. (2015). The Arabidopsis transcription factor BRASSINOSTEROID INSENSITIVE1-ETHYL METHANESULFONATE-SUPPRESSOR1 is a direct substrate of MITOGEN-ACTIVATED PROTEIN KINASE6 and regulates immunity. Plant Physiol. 167, 1076–1086. 10.1104/pp.114.250985

Lacaze A. and Joly D.L. (2020) Structural specificity in plant-filamentous pathogen interactions. Mol Plant Pathol. 21, 1513–1525. doi: 10.1111/mpp.12983

Lefebvre B., Timmers T., Mbengue M., Moreau S., Hervé C., Tóth K., Bittencourt-Silvestre J., Klaus D., Deslandes L., Godiard L., Murray J.D., Udvardi M.K., Raffaele S., Mongrand S., Cullimore J., Gamas P., Niebel A. and Ott T.(2010) A remorin protein interacts with symbiotic receptors and regulates bacterial infection. Proc Natl Acad Sci USA. 10, 2343–8. doi: 10.1073/pnas.

Li, L., Yu, X., Thompson, A., Guo, M., Yoshida, S., Asami, T., Chory, J., and Yin, Y. (2009). Arabidopsis MYB30 is a direct target of BES1 and cooperates with BES1 to regulate brassinosteroid-induced gene expression. Plant Journal. 58, 275–286. 10.1111/j.1365-313X.2008.03778.x

Li, Z., Ou, Y., Zhang, Z., Li, J., & He, Y. (2018). Brassinosteroid signaling recruits histone 3 lysine-27 demethylation activity to FLOWERING LOCUS C chromatin to inhibit the floral transition in Arabidopsis. Molecular Plant. 11, 1135–1146. 10.1016/j.molp.2018.06.007

Li Z. and He Y.(2020) Roles of Brassinosteroids in Plant Reproduction. Int J Mol Sci. 2, 872. doi: 10.3390/ijms21030872.

Liang, T., Mei, S., Shi, C., Yang, Y., Peng, Y., Ma, L., Wang, F., Li, X., Huang, X., Yin, Y., & Liu, H. (2018). UVR8 Interacts with BES1 and BIM1 to Regulate Transcription and Photomorphogenesis in Arabidopsis. Developmental Cell, 44, 512–523. 10.1016/j.devcel.2017.12.028

Lin, W., Ma, X., Shan, L., and He, P. (2013) Big roles of small kinases: the complex functions of receptor-like cytoplasmic kinases in plant immunity and development. J. Integr. Plant Biol. 55, 1188–1197

Lozano-Durán, R., Macho, A. P., Boutrot, F., Segonzac, C., Somssich, I. E., and Zipfel, C. (2013). The transcriptional regulator BZR1 mediates trade-off between plant innate immunity and growth. ELife 2, 1–15. 10.7554/eLife.00983

Malinovsky, F. G., Batoux, M., Schwessinger, B., Youn, J. H., Stransfeld, L., Win, J., Kim, S. K., & Zipfel, C. (2014). Antagonistic regulation of growth and immunity by the arabidopsis basic helix-loop-helix transcription factor HOMOLOG OF BRASSINOSTEROID ENHANCED EXPRESSION2 INTERACTING WITH INCREASED LEAF INCLINATION1 BINDING bHLH1. Plant Physiology, 164, 1443–1455. 10.1104/pp.113.234625

McLellan H., Harvey S.E., Steinbrenner J., Armstrong M.R., He Q., Clewes R., Pritchard L., Wang W., Wang S., Nussbaumer T., Dohai B., Luo Q., Kumari P., Duan H., Roberts A., Boevink P.C., Neumann C., Champouret N., Hein I., Falter-Braun P., Beynon J., Denby K. and Birch P.R.J. (2022) Exploiting breakdown in nonhost effector-target interactions to boost host disease resistance. Proc Natl Acad Sci USA. 119:e2114064119. doi: 10.1073/pnas.2114064119.

Meisrimler C.N., Pelgrom A.J.E., Oud B., Out S. and Van den Ackerveken G.(2019) Multiple downy mildew effectors target the stress-related NAC transcription factor LsNAC069 in lettuce. Plant Journal. 99, 1098–1115. doi: 10.1111/tpj.14383.

Miller M.J., Barrett-Wilt G.A., Hua Z. and Vierstra R.D. (2010) Proteomic analyses identify a diverse array of nuclear processes affected by small ubiquitin-like modifier conjugation in Arabidopsis. Proc Natl Acad Sci USA. 107,16512–7. doi: 10.1073/pnas.1004181107.

Mori, K., Lemaire-Chamley, M., Jorly, J., Carrari, F., Conte, M., Asamizu, E., and Rothan, C. (2021). The conserved brassinosteroid-related transcription factor BIM1a negatively regulates fruit growth in tomato. J of Exp Botany. 72, 1181–1197. doi:10.1093/jxb/eraa495

Mosher S and Kemmerling B. (2013) PSKR1 and PSY1R-mediated regulation of plant defense responses. Plant Signal Behav. 8(5):e24119. doi: 10.4161/psb.24119.

Mukhtar M.S., Carvunis A.R., Dreze M., Epple P., Steinbrenner J., Moore J., Tasan M., Galli M., Hao T., Nishimura M.T., Pevzner S.J., Donovan S.E., Ghamsari L., Santhanam B., Romero V., Poulin M.M., Gebreab F., Gutierrez B.J., Tam S., Monachello D., Boxem M., Harbort C.J., McDonald N., Gai L., Chen H., He Y.; European Union Effectoromics Consortium; Vandenhaute J., Roth F.P., Hill D.E., Ecker J.R., Vidal M., Beynon J., Braun P. and Dangl JL.(2011) Independently evolved virulence effectors converge onto hubs in a plant immune system network. Science. 333, 596–601. doi: 10.1126/science.1203659.

Navarro, L., Dunoyer, P., Jay, F., Arnold, B., Dharmasiri, N., Estelle, M. and Jones, J. D. (2006). A plant miRNA contributes to antibacterial resistance by repressing auxin signaling. Science, 312, 436–439. DOI:10.1126/science.1126088

Ngou B.P.M., Jones J.D.G. and Ding P. (2022) Plant immune networks. Trends Plant Sci. 27, 255–273. doi: 10.1016/j.tplants.2021.08.012.

Nota F, Cambiagno DA, Ribone P, Alvarez ME. (2015 ). Expression and function of AtMBD4L, the single gene encoding the nuclear DNA glycosylase MBD4L in Arabidopsis. Plant Sci. Jun;235:122–9.

Oh, E., Zhu, J. Y., Bai, M. Y., Arenhart, R. A., Sun, Y., & Wang, Z. Y. (2014). Cell elongation is regulated through a central circuit of interacting transcription factors in the Arabidopsis hypocotyl. ELife, 1–19. 10.7554/eLife.03031

Ohad N., Shichrur K. and Yalovsky S. (2007) The analysis of protein-protein interactions in plants by bimolecular fluorescence complementation. Plant Physiol. 145,1090–1099. doi: 10.1104/pp.107.107284.

Ortiz-Morea F.A., He P., Shan L. and Russinova E. (2020) It takes two to tango - molecular links between plant immunity and brassinosteroid signalling. J Cell Sci. 133, jcs246728. doi: 10.1242/jcs.246728.

Paik I., Kathare P.K., Kim J.I. and Huq E.(2017) Expanding Roles of PIFs in Signal Integration from Multiple Processes. Mol Plant. 10, 1035–1046. doi: 10.1016/j.molp.2017.07.002.

Ramakers, C., Ruijter, J. M., Lekanne Deprez, R. H., and Moorman, A. F. M. (2003). Assumption-free analysis of quantitative real-time polymerase chain reaction (PCR) data. Neuroscience Letters, 339, 62–66. 10.1016/S0304-3940(02)01423-4

Rehmany, A. P., Gordon, A., Rose, L. E., Allen, R. L., Armstrong, M. R., Whisson, S. C., Kamoun, S., Tyler, B. M., Birch, P. R. J., and Beynon, J. L. (2005). Differential recognition of highly divergent downy mildew avirulence gene alleles by RPP1 resistance genes from two Arabidopsis lines. Plant Cell, 17(6), 1839–1850. 10.1105/tpc.105.031807

Robert-Seilaniantz A., Grant M., Jones J.D. (2011) Hormone crosstalk in plant disease and defense: more than just jasmonate-salicylate antagonism. Annu Rev Phytopathol 2011, 49:317–343.

Romanowski A., Furniss J.J., Hussain E. and Halliday K.J. (2021) Phytochrome regulates cellular response plasticity and the basic molecular machinery of leaf development. Plant Physiol. 186, 1220–1239. doi: 10.1093/plphys/kiab112.

Salazar-Sarmiento, Felipe. 2010. Functional and molecular characterization of STH1: similarities and differences to the homolog STO. PhD thesis, Freiburg University, Freiburg,Germany. https://freidok.uni-freiburg.de/data/7868.

Sperschneider J. and Dodds P.N. (2022) EffectorP 3.0: Prediction of Apoplastic and Cytoplasmic Effectors in Fungi and Oomycetes. Mol Plant Microbe Interact. 35:146–156. doi: 10.1094/MPMI-08-21-0201-R.

Stam R., Motion G.B., Martinez-Heredia V., Boevink P.C. and Huitema E. (2021) A Conserved Oomycete CRN Effector Targets Tomato TCP14-2 to Enhance Virulence. Mol Plant Microbe Interact. 34, 309–318. doi: 10.1094/MPMI-06-20-0172-R.

Sun, Y., Fan, X., Cao, D., Tang, W., He, K., Zhu, J., He, J., Bai, M., Zhu, S., Oh, E., Patil, S., Kim, T., Ji, H., Wong, W. H., Rhee, S. Y., and Wang, Z. (2010). Article Integration of Brassinosteroid Signal Transduction with the Transcription Network for Plant Growth Regulation in Arabidopsis. Developmental Cell 19, 765–777. 10.1016/j.devcel.2010.10.010

Tör, M., Wood, T., Webb, A., Göl, D., and McDowell, J. M. (2023). Recent developments in plant-downy mildew interactions. Seminars in Cell and Developmental Biology, 10.1016/j.semcdb.2023.01.010

Toruño T.Y., Stergiopoulos I. and Coaker G.(2016) Plant-Pathogen Effectors: Cellular Probes Interfering with Plant Defenses in Spatial and Temporal Manners. Annu Rev Phytopathol. 54, 419–41. doi: 10.1146/annurev-phyto-080615-100204.

Turnbull, D., Yang, L., Naqvi, S., Breen, S., Welsh, L., Stephens, J., Morris, J., Boevink, P. C., Hedley, P. E., Zhan, J., Birch, P. R. J., and Gilroy, E. M. (2017). RXLR effector AVR2 up-regulates a brassinosteroid-responsive bHLH transcription factor to suppress immunity. Plant Physiology, 174, 356–369. 10.1104/pp.16.01804

Turnbull D, Wang H, Breen S, Malec M, Naqvi S, Yang L, Welsh L, Hemsley P, Zhendong T, Brunner F, Gilroy EM and Birch PRJ. (2019) AVR2 Targets BSL Family Members, Which Act as Susceptibility Factors to Suppress Host Immunity. Plant Physiol. 180:571–581. doi: 10.1104/pp.18.01143.

Van Butselaar, T. and Van den Ackerveken, G. (2020). Salicylic Acid Steers the Growth–Immunity Tradeoff. Trends in Plant Science, 25, 566–576. 10.1016/j.tplants.2020.02.002

Vernon L.P. (1960) Spectrophotometric Determination of Chlorophylls and Pheophytins in Plant Extracts. Analytical Chemistry. 32 (9), 1144–1150. DOI: 10.1021/ac60165a029

Verwoerd, T.C., Dekker, B.M.M. and Hoekama, A. (1989) A small-scale procedure for the rapid isolation of plant RNAs. Nucleic Acids Research, 17(6), 2362. Available from: 10.1093/nar/17.6.2362

Wang, D., Pajerowska-Mukhtar, K., Culler, A. H., & Dong, X. (2007). Salicylic Acid Inhibits Pathogen Growth in Plants through Repression of the Auxin Signaling Pathway. Current Biology. 17, 1784–1790. 10.1016/j.cub.2007.09.025

Wang H., Wang S., Wang W., Xu L., Welsh L.R.J., Gierlinski M., Whisson S.C., Hemsley P.A., Boevink P.C. and Birch P.R.J. (2023) Uptake of oomycete RXLR effectors into host cells by clathrin-mediated endocytosis. Plant Cell. 35, 2504–2526. doi: 10.1093/plcell/koad069.

Wang W., Bai M.Y. and Wang Z.Y. (2014). The brassinosteroid signaling network-a paradigm of signal integration. Current Opinion in Plant Biology 21c: 147–153.

Wang, W., Lu, X., Li, L., Lian, H., Mao, Z., Xu, P., Guo, T., Xu, F., Du, S., Cao, X., Wang, S., Shen, H., & Yang, H. Q. (2018). Photoexcited CRYPTOCHROME1 interacts with dephosphorylated bes1 to regulate brassinosteroid signaling and photomorphogenesis in arabidopsis. Plant Cell, 30(9), 1989–2005. 10.1105/tpc.17.00994

Wang, W. and Wang, Z.Y.(2014) At the intersection of plant growth and immunity. Cell Host Microbe 2014, 15, 400–402.

Wang Y., Tyler B.M. and Wang Y. (2019) Defense and Counterdefense During Plant-Pathogenic Oomycete Infection. Annu Rev Microbiol. 73, 667–696. doi: 10.1146/annurev-micro-020518-120022.

Weßling, R., Epple, P., Altmann, S., He, Y., Yang, L., Henz, S. R., McDonald, N., Wiley, K., Bader, K. C., Gläßer, C., Mukhtar, M. S., Haigis, S., Ghamsari, L., Stephens, A. E., Ecker, J. R., Vidal, M., Jones, J. D. G., Mayer, K. F. X., Ver Loren Van Themaat, E., … Braun, P. (2014). Convergent targeting of a common host protein-network by pathogen effectors from three kingdoms of life. Cell Host and Microbe, 16, 364–375. 10.1016/j.chom.2014.08.004

Win, J., Chaparro-Garcia, A., Belhaj, K., Saunders, D. G. O., Yoshida, K., Dong, S., Schornack, S., Zipfel, C., Robatzek, S., Hogenhout, S. A., & Kamoun, S. (2012). Effector biology of plant-associated organisms: Concepts and perspectives. Cold Spring Harbor Symposia on Quantitative Biology, 77, 235–247. 10.1101/sqb.2012.77.015933

Wirthmueller L., Roth C., Fabro G., Caillaud M.C., Rallapalli G., Asai S., Sklenar J., Jones A.M., Wiermer M., Jones J.D. and Banfield M.J. (2015) Probing formation of cargo/importin-α transport complexes in plant cells using a pathogen effector. Plant J. 81, 40–52. doi: 10.1111/tpj.12691

Wirthmueller L., Asai S., Rallapalli G., Sklenar J., Fabro G., Kim D.S., Lintermann R., Jaspers P., Wrzaczek M., Kangasjärvi J., MacLean D., Menke F.L.H, Banfield MJ and Jones JDG. (2018) Arabidopsis downy mildew effector HaRxL106 suppresses plant immunity by binding to RADICAL-INDUCED CELL DEATH1. New Phytol. 220, 232–248. doi: 10.1111/nph.15277.

Xing, S., Quodt, V., Chandler, J., Höhmann, S., Berndtgen, R., & Huijser, P. (2013). SPL8 acts together with the brassinosteroid-signaling component BIM1 in controlling Arabidopsis thaliana male fertility. Plants. 2, 416–428. 10.3390/plants2030416

Xu, P., Fang, S., Chen, H., and Cai, W. (2020). The brassinosteroid-responsive xyloglucan endotransglucosylase/hydrolase 19 (XTH19) and XTH23 genes are involved in lateral root development under salt stress in Arabidopsis. Plant Journal, 104, 59–75. 10.1111/tpj.14905

Yang L., Teixeira P.J., Biswas S., Finkel O.M., He Y., Salas-Gonzalez I., English M.E., Epple P., Mieczkowski P. and Dangl J.L.(2017) Pseudomonas syringae Type III Effector HopBB1 Promotes Host Transcriptional Repressor Degradation to Regulate Phytohormone Responses and Virulence. Cell Host Microbe. 21,156–168. doi: 10.1016/j.chom.2017.01.003.

Yin, Y., Wang, Z. Y., Mora-Garcia, S., Li, J., Yoshida, S., Asami, T., and Chory, J. (2002). BES1 accumulates in the nucleus in response to brassinosteroids to regulate gene expression and promote stem elongation. Cell, 109, 181–191. 10.1016/S0092-8674(02)00721-3

Yin, Y., Vafeados, D., Tao, Y., Yoshida, S., Asami, T., and Chory, J. (2005). A new class of transcription factors mediates brassinosteroid-regulated gene expression in Arabidopsis. Cell, 120, 249–259. 10.1016/j.cell.2004.11.044

Yu, X., Li, L., Zola, J., Aluru, M., Ye, H., Foudree, A., Guo, H., Anderson, S., Aluru, S., Liu, P., Rodermel, S., and Yin, Y. (2011). A brassinosteroid transcriptional network revealed by genome-wide identification of BESI target genes in Arabidopsis thaliana. Plant Journal, 65(4), 634–646. 10.1111/j.1365-313X.2010.04449.

Yuan J. and He S.Y. (1996) The Pseudomonas syringae Hrp regulation and secretion system controls the production and secretion of multiple extracellular proteins. J. Bacteriol. 178:6399–6402.

Zhang L.Y., Bai M.Y., Wu J., Zhu J.Y., Wang H., Zhang Z., Wang W., Sun Y., Zhao J., Sun X., Yang H., Xu Y., Kim S.H., Fujioka S., Lin W.H., Chong K., Lu T. and Wang Z.Y. (2009) Antagonistic HLH/bHLH transcription factors mediate brassinosteroid regulation of cell elongation and plant development in rice and Arabidopsis. Plant Cell. 21, 3767–80. doi: 10.1105/tpc.109.070441.

Zhu J.Y., Sae-Seaw J. and Wang Z.Y. (2013) Brassinosteroid signalling. Development 140: 1615–1620.

Zhu Z., Xu F., Zhang Y., Cheng Y.T., Wiermer M., Li X. and Zhag Y (2010). Arabidopsis resistance protein SNC1 activates immune responses through association with a transcriptional corepressor. Proc Natl Acad Sciences USA. 107, 13960–13965. 10.1073/pnas.1002828107

Züst T. and Agrawal A.A. (2017) Trade-Offs Between Plant Growth and Defense Against Insect Herbivory: An Emerging Mechanistic Synthesis. Annu Rev Plant Biol. 68, 513–534. doi: 10.1146/annurev-arplant-042916-040856.

Zuo J., Niu Q.W. and Chua N.H. (2000) Technical advance: An estrogen receptor-based transactivator XVE mediates highly inducible gene expression in transgenic plants. Plant J. 24:265–73. doi: 10.1046/j.1365-313x.2000.00868.x.

