## Supplementary Figures for "Downy mildew effector HaRxL106 interacts with the transcription factor BIM1 altering plant growth, BR signaling and susceptibility to pathogens"

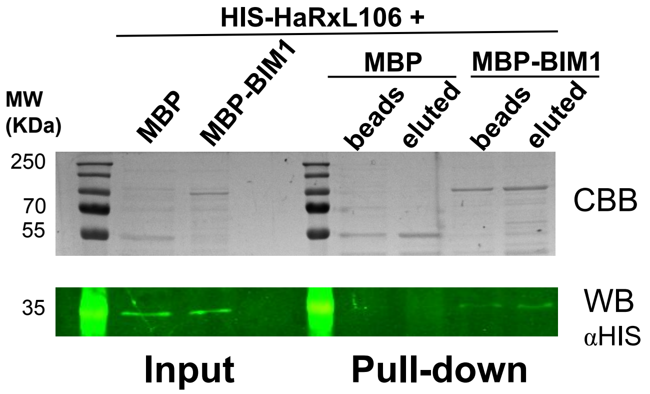


**Supplementary Figure S1: HaRxL106 physically interacts with BIM1.**
*In vitro* pull-down assay performed with MBP-BIM1 and HIS-HaRxL106. Recombinant MBP-BIM1 or MBP alone (negative control) produced in *E. coli* were bound to an amylose resin and then mixed with HIS-HaRxL106 previously purified from a crude extract of soluble *E. coli* proteins via a Ni-NTA column. Input fractions as well as post-washed amylose beads (beads) and maltose eluted (eluted) fractions were collected and separated by SDS-PAGE. Identical gels were used for coomassie blue staining (CBB) or western blot (WB), revealed with primary anti-HIS antibody.

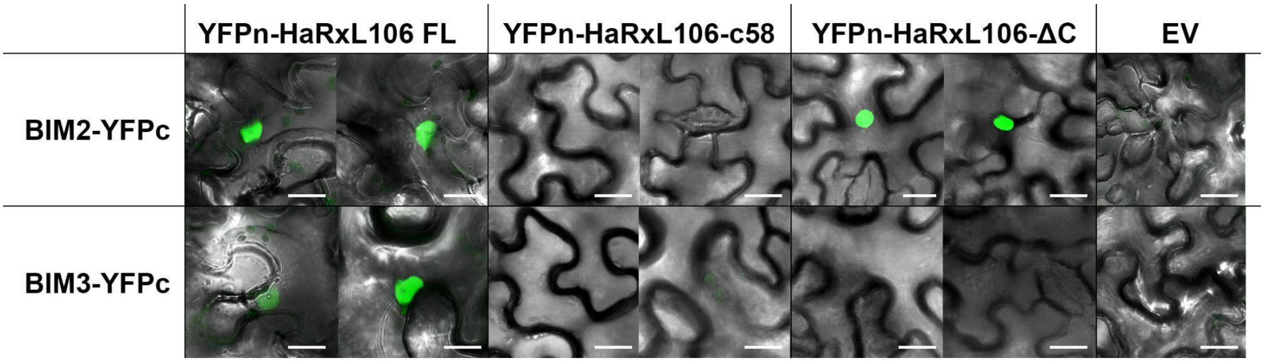


**Supplementary Figure S2: HaRxL106 interacts with BIM2 and BIM3.** Confocal microscopy images representative of the results obtained when a Bi-Molecular fluorescence complementation assay (BiFC) was performed in *N. benthamiana* leaves co-infiltrated with each pair of the indicated constructs. Scale bars represent 20 μm.

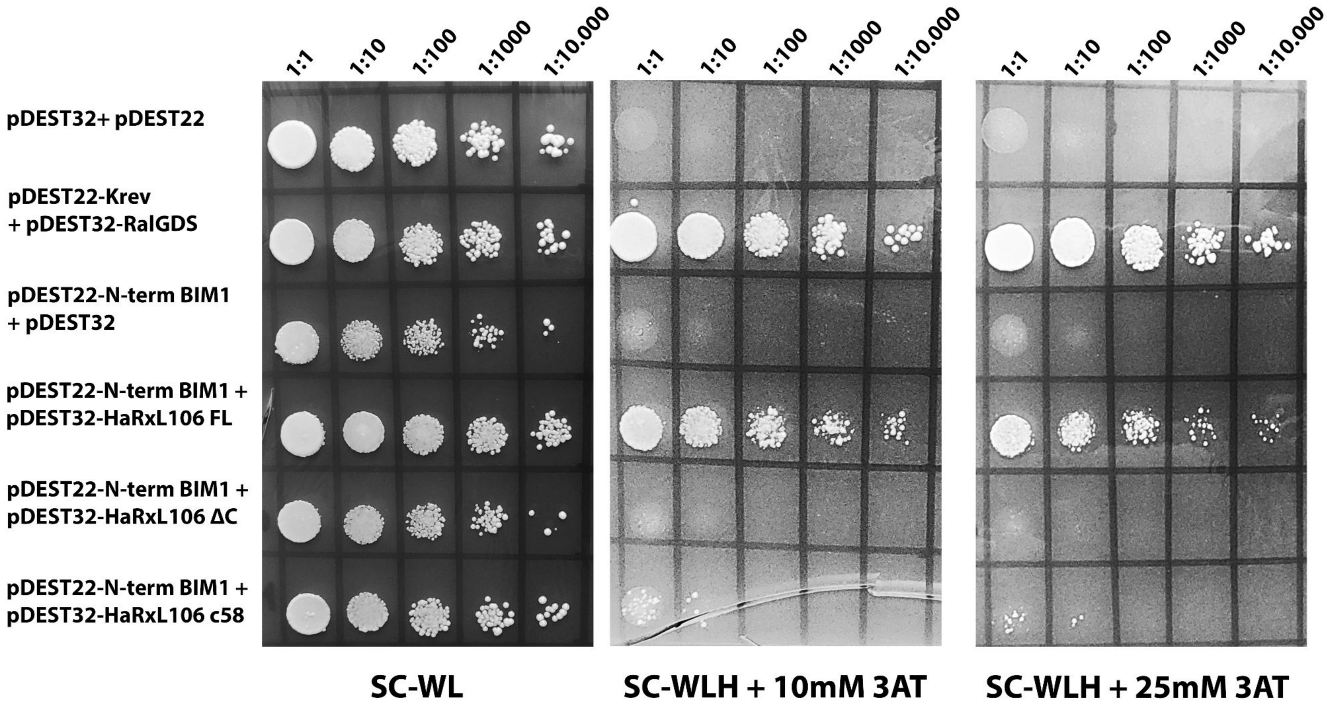


**Supplementary Figure S3**: **The N-terminal domain of BIM1 interacts with full length HaRxL106, but not with the effector C-terminal domain.** Yeast-two-Hybrid assay performed with the N-terminal domain of BIM1 (N-term BIM1) cloned into pDEST22 prey vector. The effector HaRxL106 either the full length (FL) version as well as the carboxyl (HaRXL106-c58) and amino-terminal (HaRxL106-ΔC) domains were cloned into pDEST32 bait vector. SC-WL: medium Synthetic Complete for yeasts without Tryptophan and Leucine. SC-WLH + 10/25 mM 3AT: SC-WL without Histidine, supplemented with 10 or 25 mM 3-Aminotriazole. pDEST32-Krev1+pDEST22-RalGDS: positive interaction control.

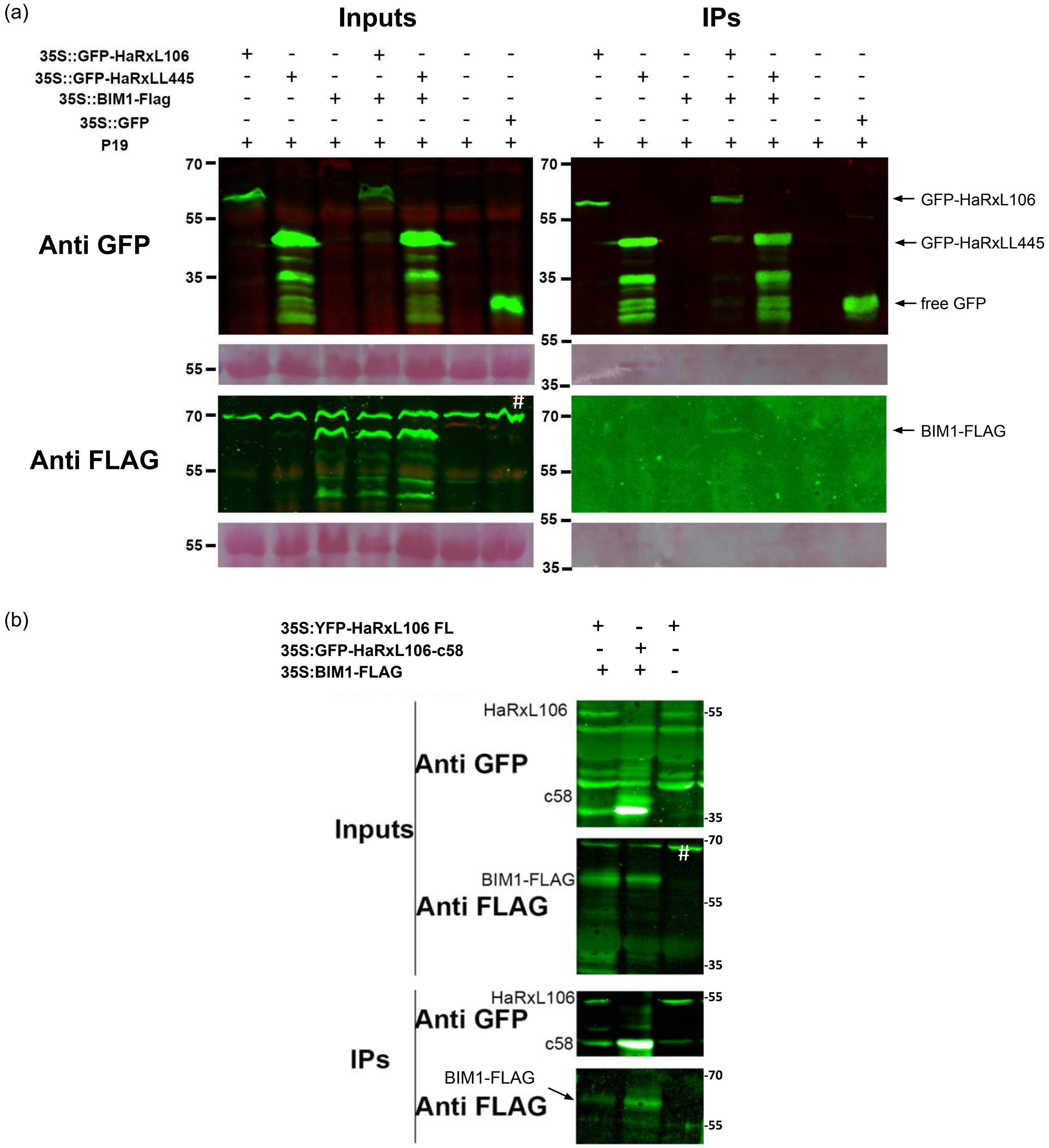


**Supplementary Figure S4**: **GFP-HaRxL106 full length and GFP-HaRxL106-c58 co-immunoprecipitate with BIM1-FLAG in *N. benthamiana*. a.** Left panels: electrophoretic runs of total protein extracts (inputs), revealed with anti-GFP (upper panel) or anti-FLAG antibodies (lower panel). Right panels: proteins immunoprecipitated (IPs) with Magne-Halo-anti-GFP, revealed with anti-GFP (upper panel) or anti-FLAG antibodies (lower panel). (#) Unspecific band observed with anti-FLAG antibody. **b.** Inputs: total protein extracts, revealed with anti-GFP (upper panel) or anti-FLAG antibodies (lower panel). IPs: proteins immunoprecipitated with anti-GFP antibody (Magne-Halo-antiGFP), revealed with anti-GFP antibody (upper panel) or with anti-FLAG antibody (lower panel).

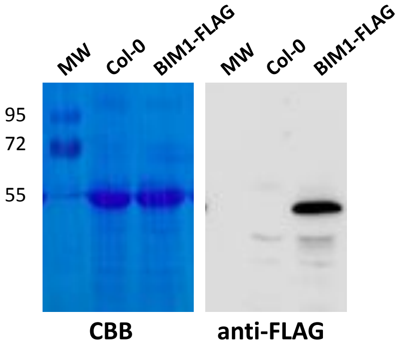


**Supplementary Figure S5: BIM1-FLAG fusion protein is expressed in Arabidopsis seedlings.** Left panel: coomassie brilliant blue staining of RUBisCO large subunit as loading control of the western blot shown on the right side. Molecular weight of the marker is shown with numbers (kDa).

Right panel: Western blot showing BIM1-FLAG detection by anti-FLAG antibody in a total protein extract from BIM1-FLAG *A. thaliana* transgenic seedlings. Col-0 protein extract is used as negative control to reveal unspecific bands.

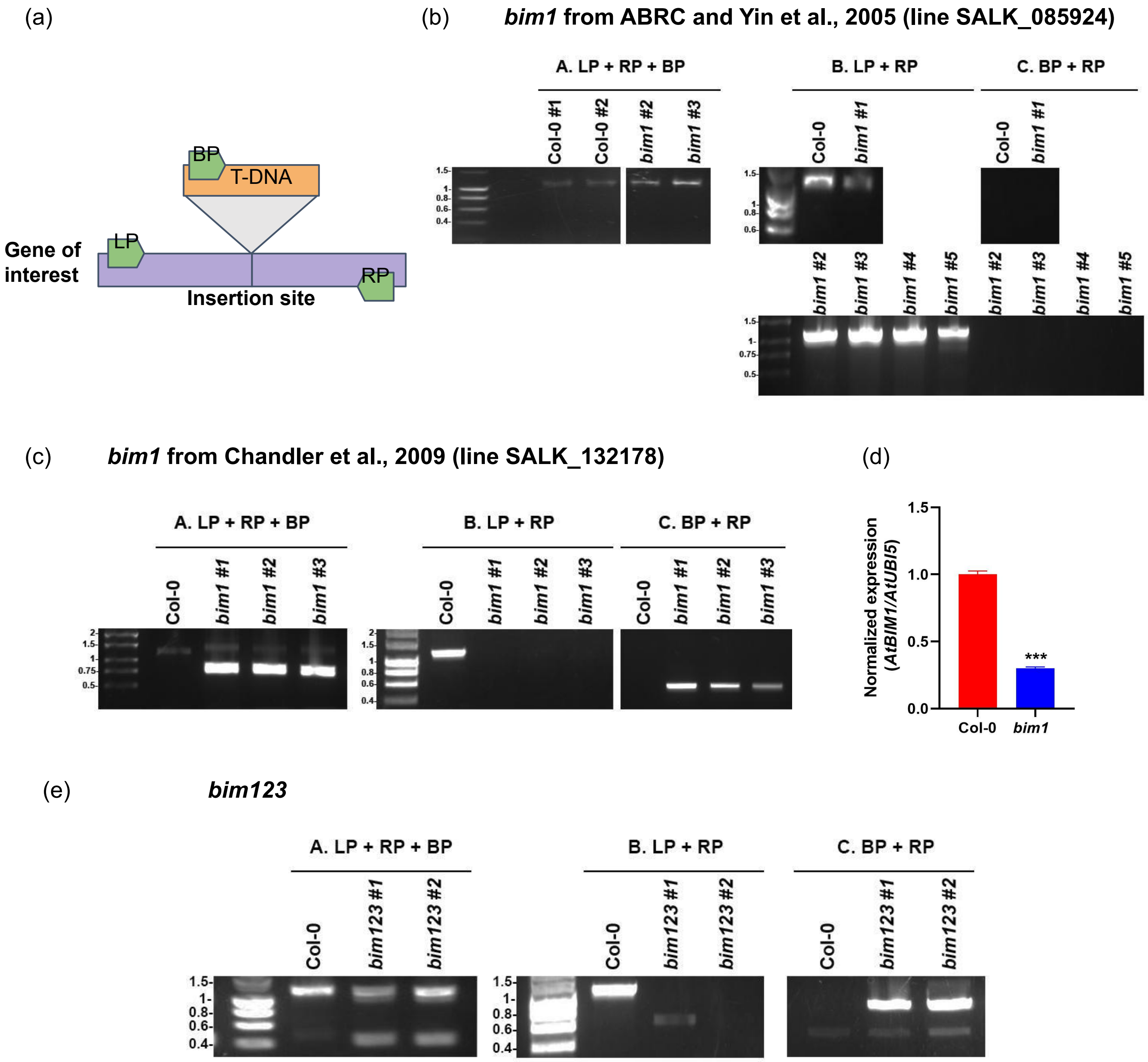


**Supplementary Figure S6: SALK_132178 mutant plants are knockdown mutants for BIM1. a**. Diagram showing the regions on which the primers are expected to hybridize. LP: left primer, RP: right primer, BP: primer on the T-DNA, all listed in Supplementary Table 2. **b**. Electrophoretic runs in agarose gels of PCRs performed with different primer combination (A, B and C) on genomic DNA of Col-0 and five plants of the line SALK_085924. **c**. Electrophoretic runs in agarose gels of PCRs performed with the mentioned primer combinations on genomic DNA of Col-0 and three plants of the line SALK_132178. **d**. Expression of BIM1 transcript normalized to the reference gene (*AtUBI5*) in Col-0 and line SALK_132178 (*bim1)* seedlings. Asterisks indicate significant differences according to bilateral t-test with p<0.001. **e**. **Screening of *bim123* triple mutants.** Electrophoretic runs in agarose gels of PCRs performed with the mentioned primer combinations on genomic DNA of Col-0 and two plants of the *bim123* triple mutant. The primers used here were designed to screen the insertion at the 5' end of BIM1 (SALK_085924 line).

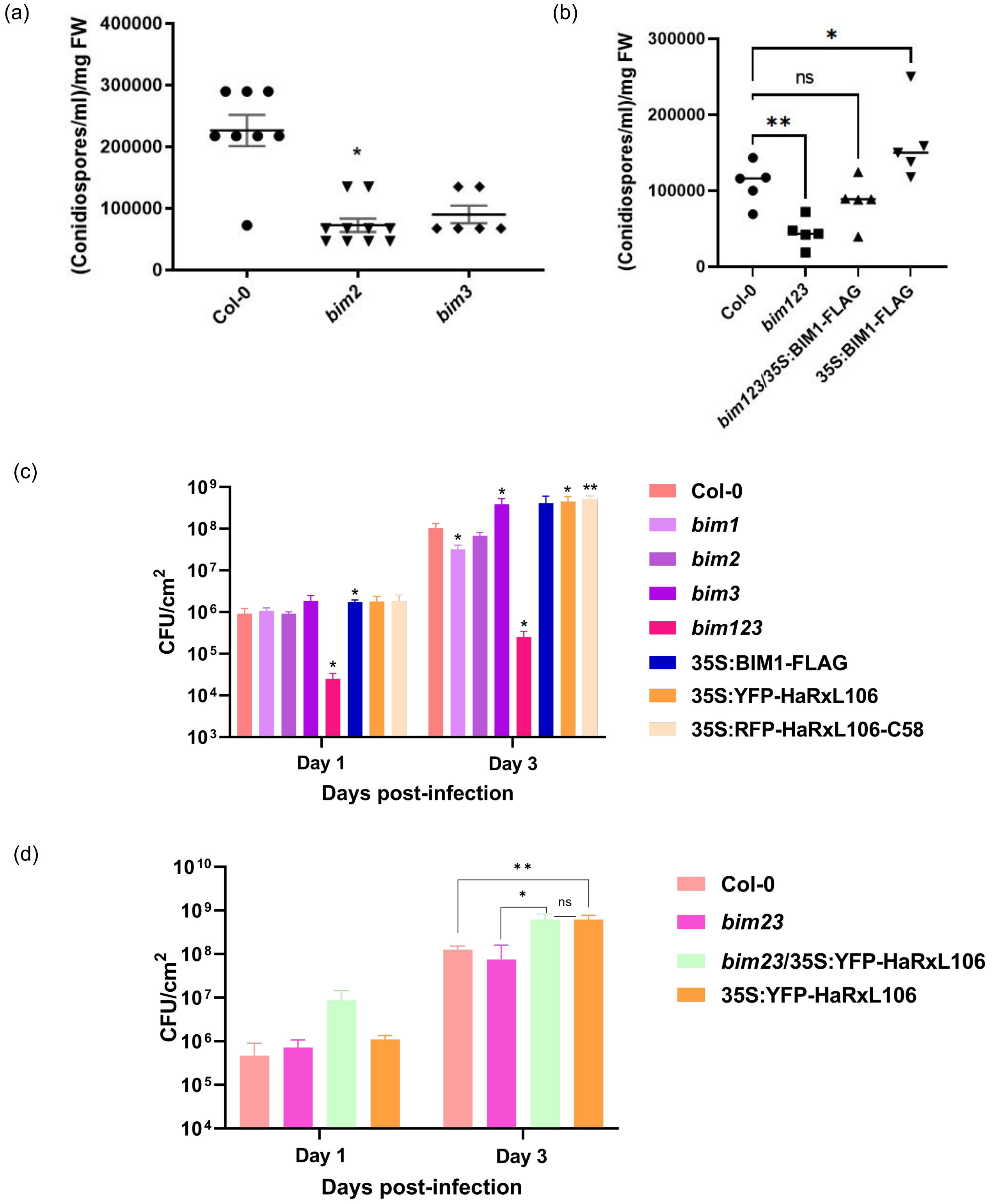


**Supplementary Figure S7: Susceptibility of single (*bim1, bim2 and bim3*), double (*bim23*) and triple (*bim123*) mutants to *Hyaloperonospora arabidopsidis* (Hpa) NoCo2 and *Pseudomonas syringae* pv. tomato DC3000 (Pst). a.** Hpa infection assay using $10^5$ spores/ml in 7-day-old *A. thaliana* seedlings of the indicated genotypes. Asterisk indicate significant differences compared to wild type (Col-0) plants according to non-parametric ANOVA (Kruskal Wallis) with p<0.05. **b**. Hpa infection assay using $10^7$ spores/ml in 15-day-old *A. thaliana* seedlings of the indicated genotypes. Asterisks indicate significant differences compared to wild type (Col-0) plants according to a parametric ANOVA (uncorrected Fisher's LSD post hoc test) with p<0.05 (*) or p <0.01 (**). ns: not statistically significant. **c-d.** Bacterial growth curves assays indicating that BIM2 and BIM3 are not necessary for the increased susceptibility to Pst of HaRxL106 overexpressing plants. Colony-forming units per square centimeter of leaf ($CFU/cm^2$) were determined at 1 and 3 days syringe inoculation of the bacteria in the genotypes mentioned. Significant differences with the wild type (Col-0) or between the genotypes indicated according to a t-test for averaged differences with p<0.05 (*) and p <0.01 (**) are indicated. ns: not statistically significant.

(a)

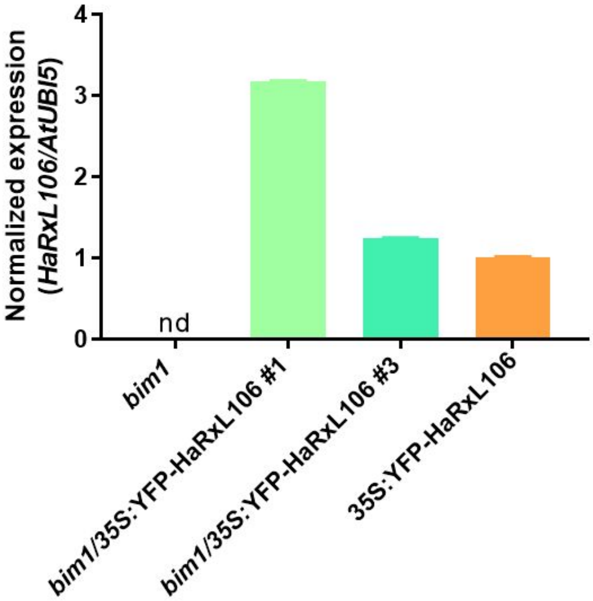


(b)

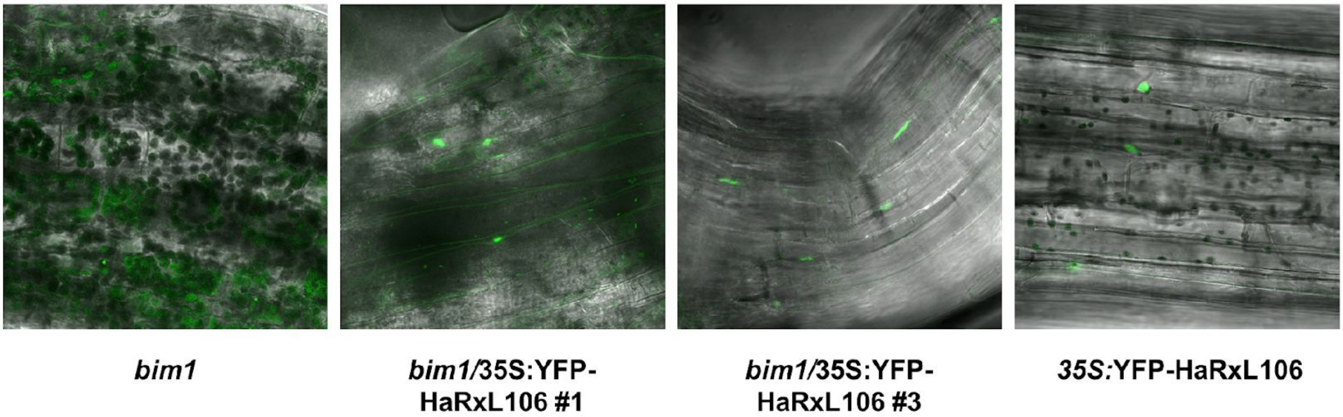


**Supplementary Figure S8: *bim1*/35S:YFP-HaRxL106 lines express the HaRxL106 transcript and GFP-HaRxL106 fusion protein. a.** Expression of 156 pb of the N-terminal of HaRxL106 transgene normalized to a reference gene expression (*AtUBI5*) in the specified genotypes. nd: not detected. **b.** Confocal microscopy images of hypocotyls of seedlings of the indicated genotypes. *bim1*: line SALK_132178. Fluorescent nuclei indicate the presence of the fusion protein in the predicted subcellular localization.

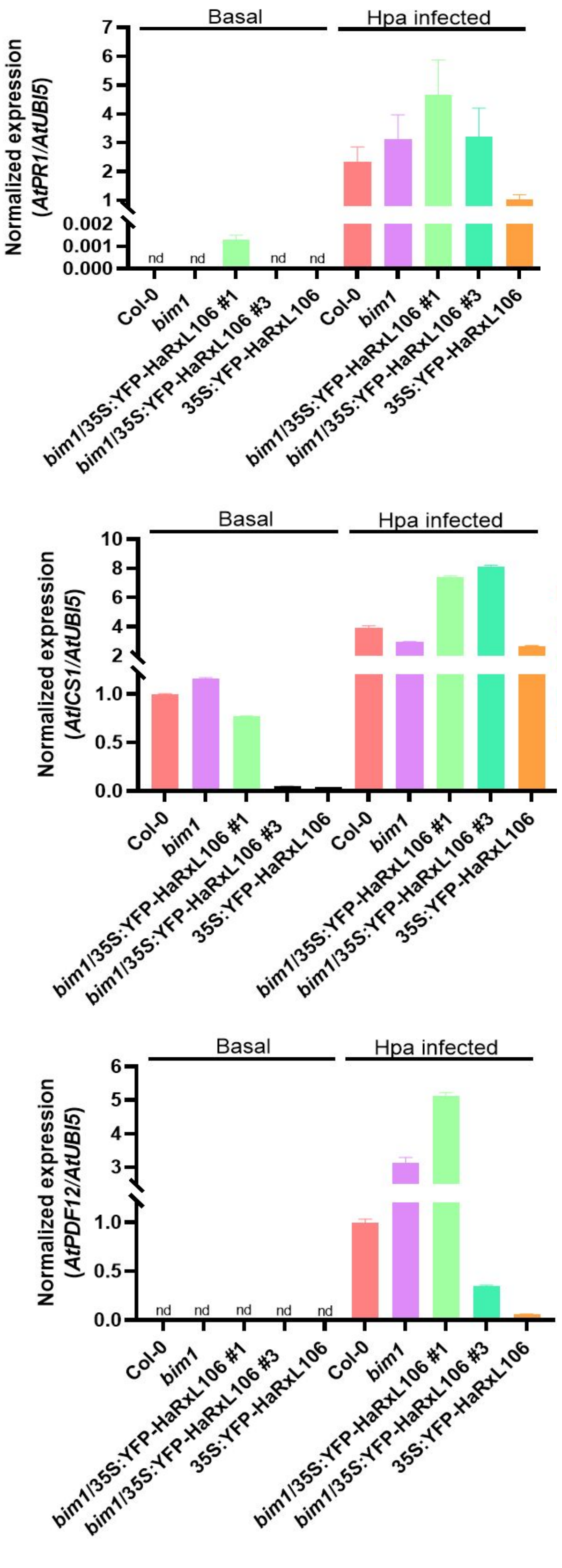


**Supplementary Figure S9: Defense gene expression in Arabidopsis seedlings of different genotypes before and after Hpa infection.** Expression of the indicated genes was determined in 10-day-old seedlings of different genotypes by qPCR with the primers listed on Supplementary table 2. nd: not detected.

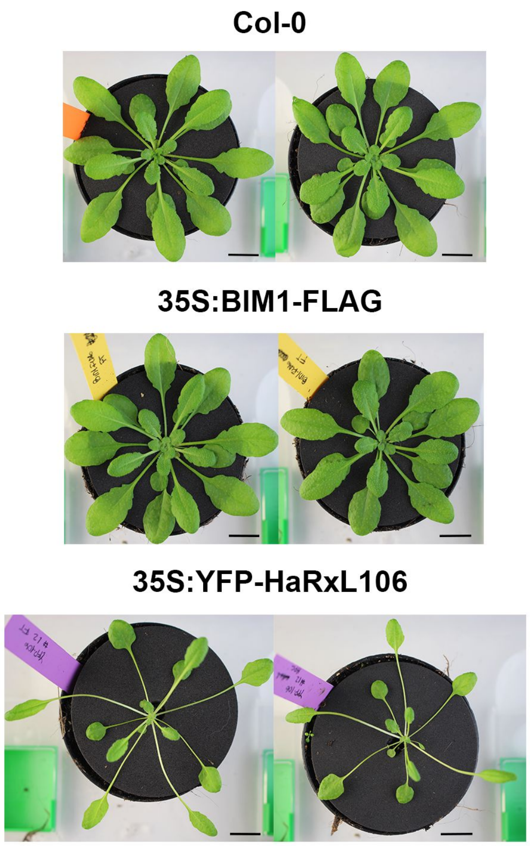


**Supplementary Figure S10: BIM1-FLAG overexpression does not phenocopies the SAS-like phenotype of HaRxL106 transgenic lines.** Phenotype of two-month-old adult plants of BIM1-FLAG compared to same age Col-0 and Col-0 expressing HaRxL106. Scale bars represent 1,5 cm.

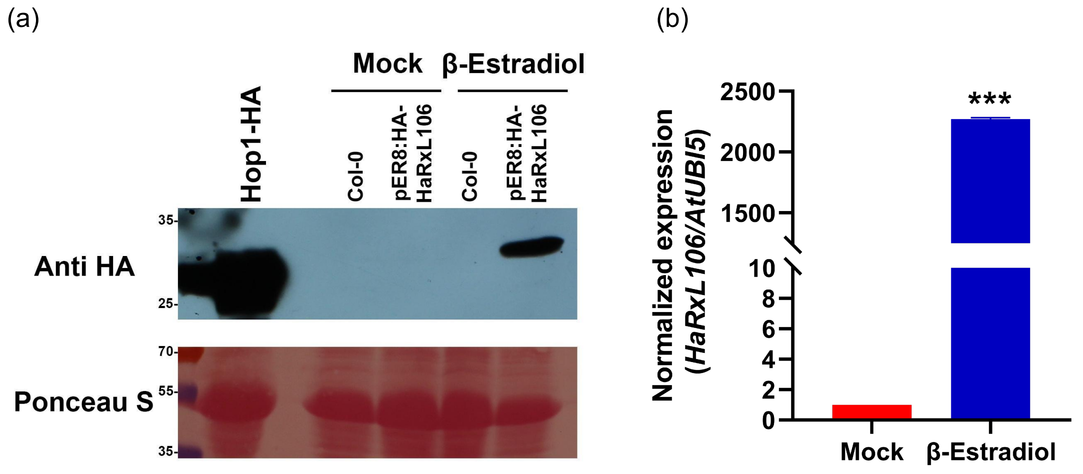


**Supplementary Figure S11: Verification that a single treatment with β-estradiol is effective to induce the expression of HaRxL106 transcript and protein in pER8:HA-HaRxL106 plants. a.** Expression of HA tagged HaRxL106 protein in seedlings of the indicated genotypes, treated with DMSO (mock) or β-Estradiol. Top panel: Total protein extracts, revealed with Anti-HA antibody. Bottom panel: Loading control of the RuBisCO large subunit protein stained with Ponceau S. **b**. Normalized expression of the HaRxL106 transcript (156bp) in response to β-estradiol treatment, 1.25 μM final concentration in MS plates**.** *** indicates statistically significant differences between mock and β-Estradiol treated pER8:HA-HaRxL106 plants according to a unpaired t-test ($p<0.05$).

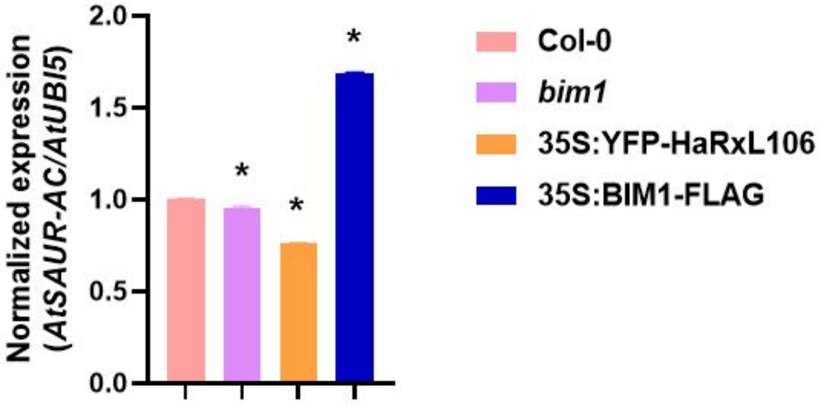


**Supplementary Figure S12: BIM1-FLAG and HaRxL106 affect the transcription of SAUR-AC gene.** Expression of SAUR-AC normalized to the reference gene (*AtUBI5*) in 10-day-old seedlings of the indicated genotypes. * indicates statistically significant differences regarding genotype Col-0, according to a one-factor ANOVA (Tukey post hoc test, $p<0.05$).

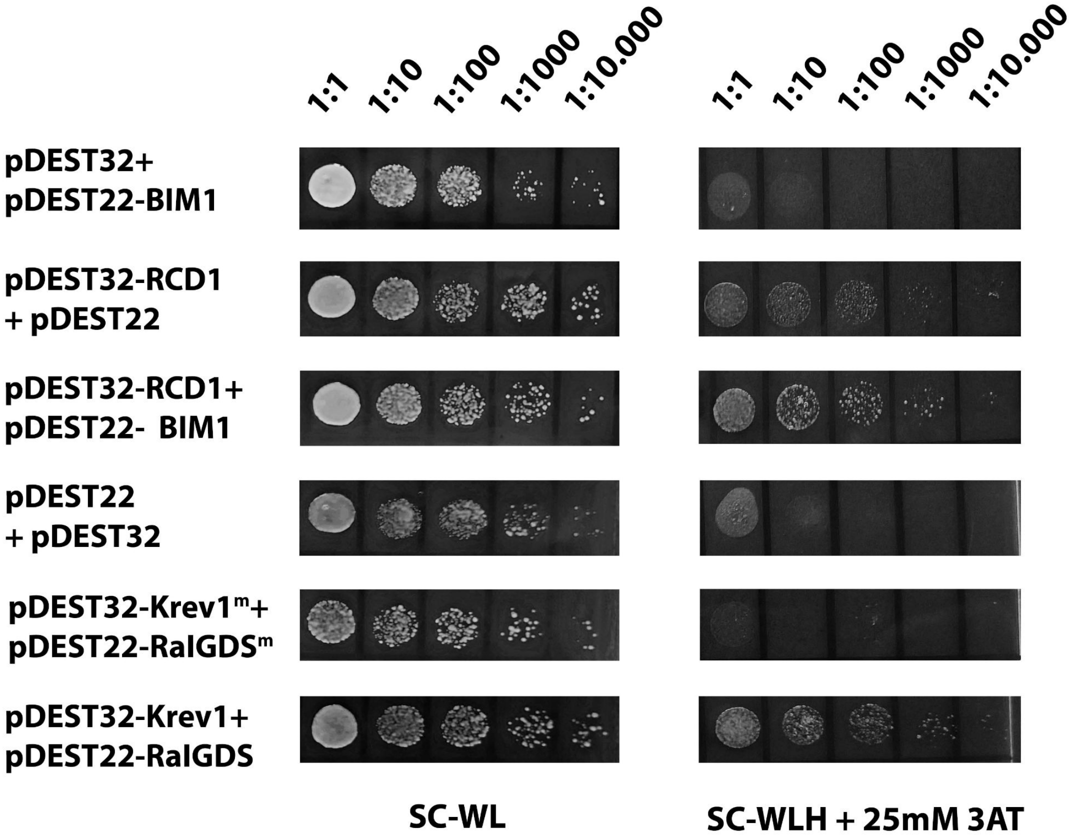


**Supplementary Figure S13: BIM1 and RCD1, the nuclear targets of HaRxL106, interact between them in Y2H.**

Y2H assay performed with BIM1 cloned into the prey vector (pDEST22) and RCD1 into the bait vector (pDEST32). BIM1 was able to interact with the transcriptional regulator RCD1. SC-WL: medium Synthetic Complete for yeasts without Tryptophan and Leucine. SC-WLH + 25 mM 3AT: SC-WL without Histidine, supplemented with 25mM 3-Aminotriazole. pDEST32-Krev1+pDEST22-RalGDS: positive interaction control. pDEST32-Krev1$^{m}$+pDEST22-RalGDS$^{m}$: negative interaction control.
