## Supplementary Tables for "Downy mildew effector HaRxL106 interacts with the transcription factor BIM1 altering plant growth, BR signaling and susceptibility to pathogens"

**Supplementary Table 1:** List of resources used in this work

| Resource | Identifier | Source |
|---|---|---|
| *Seeds* | | |
| *bim1* single mutant | N654404/SALK_132178C | Chandler et al., 2009 |
| *bim2* single mutant | N668502/SALK_074689C | ABRC |
| *bim3* single mutant | N579683/SALK_079683 | ABRC |
| *bim23* double mutant | | This work |
| *bim123* triple mutant | | Liang et al., 2018 |
| 35S:BIM1-FLAG | | Liang et al., 2018 |
| 35S:YFP-HaRxL106 | | Wirthmueller et al., 2018 |
| 35S:RFP-HaRxL106-c58 | | Wirthmueller et al., 2018 |
| 35S:GFP-HaRxL106 FL | | This work |
| 35S:GFP-HaRxL106-ΔC | | This work |
| 35S:GFP-HaRxL106-c58 | | This work |
| *bim1*/35S:YFP-HaRxL106 | | This work |
| *bim23*/35S:YFP-HaRxL106 | | This work |
| *bim123*/35S:YFP-HaRxL106 | | This work |
| 35S:BIM1-FLAG/ 35S:YFP-HaRxL106 FL | | This work |
| 35S:BIM1-FLAG/ 35S:YFP-HaRxL106-ΔC | | This work |
| 35S:BIM1-FLAG/ 35S:YFP-HaRxL106-c58 | | This work |
| pER8:HA-HaRxL106 | | G. Fabro (J.D.G. Jones Lab, TSL, Norwich, UK) |
| 35S:GFP | | Nota el al., 2015 |
| *Plasmids* | | |
| pENTRY-D-TOPO-HaRxL106 FL | | Wirthmueller et al., 2015 |

| | | |
|---|---|---|
| pENTR4-HaRxL106 FL | | Wirthmueller et al., 2018 |
| pENTRY-D-TOPO-SV40NLS-HaRxL106-ΔC | | Wirthmueller et al., 2015 |
| pENTRY-D-TOPO-HaRxL106-c58 | | Wirthmueller et al., 2015 |
| pENTRY-D-TOPO-BIM1 | | This work |
| pENTR4-BIM2 | | This work |
| pENTR4-BIM3 | | This work |
| 35S:GFP-GW (pK7WGF2) | | Ghent University |
| 35S:GW-GFP (pB7FWG2) | | Ghent University |
| BIM1-CFPc | | Liang et al., 2018 |
| BIM1-GFP | | This work |
| BIM1-FLAG | | Liang et al., 2018 |
| MBP-BIM1 | | Liang et al., 2018 |
| BIM2-YFPc | | This work |
| BIM3-YFPc | | This work |
| MOS6-YFPc | | Wirthmueller et al., 2015 |
| HIS-HaRxL106 | | Wirthmueller et al., 2015 |
| GFP-HaRxL106 | | This work |
| YFP-HaRxL106 | | Wirthmueller et al., 2018 |
| YFPn-HaRxL106 | | Wirthmueller et al., 2015 |
| YFPn-HaRxL106-ΔC | | Wirthmueller et al., 2015 |
| YFPn-HaRxL106-c58 | | This work |
| PRE5pro:LUC | | Liang et al., 2018 |
| SAUR-ACpro:LUC | | Liang et al., 2018 |
| pDEST22-BIM1 | | This work |
| pDEST22-N Term BIM1 | | This work |
| pDEST32-HaRxL106 FL | | Wirthmueller et al., 2018 |
| pDEST32-HaRxL106-ΔC | | Wirthmueller et al., 2018 |
| pDEST32-HaRxL106-c58 | | This work |

| | | |
|---|---|---|
| pDEST32-RCD1 | | Wirthmueller et al., 2018 |

**Supplementary Table 2:** List of primers used in this work

| Primer | Sequence | Used for |
|---|---|---|
| BIM1-LP (SALK_085924) | GCGTACGAGCTGCAATTAGAG | Genotypic analysis of T-DNA insertional mutant lines |
| BIM1-RP (SALK_085924) | CAACTTCAGAGCTCAATTCCG | Genotypic analysis of T-DNA insertional mutant lines |
| BIM1-LP (SALK_132178) | TGTATTGTCCGAGATTCCACC | Genotypic analysis of T-DNA insertional mutant lines |
| BIM1-RP (SALK_132178) | CACTCCTGAATTCTCCAATGC | Genotypic analysis of T-DNA insertional mutant lines |
| BIM2-LP (SALK_074689) | ATGCTTTGTGCGAATATCCTG | Genotypic analysis of T-DNA insertional mutant lines |
| BIM2-RP (SALK_074689) | CAGATTCTTCCTCAACGCTTG | Genotypic analysis of T-DNA insertional mutant lines |
| BIM3-LP (SALK_079683) | TGTTTTTGGTTGCCAAAAGTC | Genotypic analysis of T-DNA insertional mutant lines |
| BIM3-RP (SALK_079683) | ACTGTTTGAATTCGGATGCAG | Genotypic analysis of T-DNA insertional mutant lines |
| LBb1.3 SALK (primer BP) | ATTTTGCCGATTTCGGAAC | Genotypic analysis of T-DNA insertional mutant lines |
| BIM1_no_start_FW | CAAAAAAGCAGGCTCCACTGAGCTTCCTCAACCTCGTC | Cloning by Gibson of BIM1 splicing variant 5 CDS in pENTR4 for N-terminal tag |
| BIM1_stop_RV | GCTGGGTCTAGATATCCTACTGTCCCGTCTTGAGC | Cloning by Gibson of BIM1 splicing variant 5 CDS in pENTR4 for N-terminal tag |

| | | |
|---|---|---|
| BIM1-GW-FW | CACCATGGAGCTTCCTCAACC | Cloning of BIM1 splicing variant 5 in pENTR-D-TOPO for N or C-terminal tag |
| BIM1-GW-RV_no stop | CTGTCCCGTCTTGAGCCG | Cloning of BIM1 splicing variant 5 in pENTR-D-TOPO for N or C-terminal tag |
| BIM1-GW-RV_stop | CTACTGTCCCGTCTTGAGCCG | Cloning of BIM1 splicing variant 5 in pENTR-D-TOPO for N or C-terminal tag |
| BIM1 middle FW | GCAGCAACTGTGGGACAATG | Sequencing. Anneals in the central region of BIM1 CDS. |
| BIM1 middle RV | TCCACTCTATGGCTCTGGCT | Sequencing. Anneals in the central region of BIM1 CDS. |
| pENTR4_fw | CCCGCCATAAACTGCCAGG | *Sequencing. Anneals on* pENTR4 backbone. |
| pENTR4_rv | CGTTGAATATGGCTCATAACACCC | *Sequencing. Anneals on* pENTR4 backbone. |
| M13 FW | GTTGTAAAACGACGGCCAGT | Sequencing of destination vectors. |
| M13 RV | CAGGAAACAGCTATGAC | Sequencing of destination vectors. |
| ARF6_exp_FW | CAAAGTTTAGCAGCTACCACGA | qPCR |
| ARF6_exp_RV | ACGTCGTTCTCTCGGTCAAC | qPCR |
| BIM1_exp_FW | CTCTCTTCCTCTCAAGGGTCTGT | qPCR |
| BIM1_exp_RV | CTCTCACATCTGCTTTTACCCTCA | qPCR |
| BR6OX_exp_fw | TGTGGTTGGGATGATCTTGA | qPCR |
| BR6OX_exp_rv | CTCCACTGCGGTAATTCGTT | qPCR |
| DWF4_exp_fw | AGGTGGGATTCTTGGGAAAT | qPCR |

| | | |
|---|---|---|
| DWF4_exp_rv | CTTTTTGGCCTCGTCTTGAG | qPCR |
| HaRxL106_exp_fw | ACAACGGCAACGAAGAGAGAAATG | qPCR |
| HaRxL106_exp_rv | CGCTATACAAAGGAAGGCTCGAC | qPCR |
| IAA19_exp_fw | GGAAGATGGATCTTGGTTCG | qPCR |
| IAA19_exp_rv | CATCCCCCAAGGTACATCAC | qPCR |
| ICS1_exp_fw | CTGCTGTAGAGAAGGCTTTAGAGATGA | qPCR |
| ICS1_exp_rv | AGTCTCTCAGGCGTGTTTCCGAT | qPCR |
| MYB30_exp_fw | GCGAAAAAGGCTCTCTCTGA | qPCR |
| MYB30_exp_rv | TTTTTCACCCACCCTTTGAG | qPCR |
| PDF12_N_fw | TTGCTGCTTTCGACGCA | qPCR |
| PDF12_N_rv | TGTCCCACTTGGCTTCTCG | qPCR |
| PR1_exp_fw | ATGAATTTTACTGGCTATTCTC | qPCR |
| PR1_exp_rv | AGGGAAGAACAAGAGCAACTA | qPCR |
| PRE1_exp_fw | TCAAGGCAATCTTCAAGTGC | qPCR |
| PRE1_exp_rv | AGACAAACGCTCGCTCAGAT | qPCR |
| PRE5_exp_fw | TTCGAATGCTTCGAGGATCT | qPCR |
| PRE5_exp_rv | AAGAAGCTGCGACAAACGAT | qPCR |

| | | |
|---|---|---|
| SAUR-AC_exp_fw | TGAGGAGTTTCTTGGGTGCT | qPCR |
| SAUR-AC_exp_rv | TATTGTTAAGCCGCCCATTG | qPCR |
| UBI5_exp_fw | GTGGTGCTAAGAAGAGGAAGA | qPCR |
| UBI5_exp_rv | TCAAGCTTCAACTCCTTCTTT | qPCR |
| XTH19_exp_fw | TTCACGATAATCAAGGGAAAC | qPCR |
| XTH19_exp_rv | AAAGATAGAATGTTGTGACGG | qPCR |
